# Paradoxical non-catalytic kinase functions are driven by inhibitor-induced displacement of autoinhibitory domains

**DOI:** 10.1101/2025.11.06.687012

**Authors:** Viviane Reber, Sabrina Keller, Stefanie A. Loosli, Yumi Arima, Tatjana Kleele, Paola Picotti, Matthias Gstaiger

**Author notes:** Corresponding author: Matthias Gstaiger.

## Abstract

ATP competitive kinase inhibitors represent one of the largest classes of targeted anti-cancer drugs. While their primary mechanism is to block catalytic activity, they can also trigger paradoxical phenotypic effects that cannot be explained by catalytic inhibition alone. These observations point to a hidden layer of drug action that modulates non-catalytic kinase functions via changes in kinase conformation and protein-protein interactions (PPIs). Here we developed a multimodal proteomics approach combining limited proteolysis coupled mass spectrometry on affinity purified samples (AP-LiP-MS), AP-MS, and proximity labeling-MS to map inhibitor-induced conformation and PPI changes. We show that inhibitor binding causes structural rearrangements in the AIDs of all tested kinases, consistent with a transition to an open, active-like kinase conformation. These structural shifts drive distinct kinase-protein interaction changes that control non-catalytic functions: sequestration of AMPK by inhibited CAMKK2 blocks phosphorylation by other kinases, CHEK1 inhibition causes dissociation from the mitochondrial protein CLPB and leads to mitochondrial fragmentation, and structural changes in inhibited PRKCA trigger rapid relocalization to cell junctions. Thus we identify the ATP-binding site as a major organizing center of kinase conformation and interaction. Our work suggests that these on-target, off-mechanism effects are highly prevalent and provides the analytical framework to systematically characterize a frequently overlooked phenomenon highly relevant for understanding drug side effects to guide the development of novel therapeutics.

## Introduction

Most human proteins are phosphorylated by the 518 kinases^1^, representing a key mechanism to organize the dynamic functional landscape of the proteome orchestrating the majority of cellular processes. Consequently, mutations and changes in signaling context that affect kinase function are often causally linked to the development of human diseases such as cancer^2,3^. Therapeutic efforts to correct aberrant kinase activity focused on the development of compounds which bind the kinase domain (KD) and inhibit its catalytic activity. This resulted in a set of chemical probes targeting over 130 kinases^4^. Most probes targeting kinases inhibit their catalytic activity in an ATP-competitive manner (also referred to as a type I inhibitor), representing one of the largest classes of FDA approved anti-cancer drugs^5,6^.

Classical target engagement analyses typically focus on kinetic parameters of catalytic inhibition, target specificity and structural characterization of the probe-KD interaction. However, beyond the catalytic core, protein kinases contain multiple additional domains that confer diverse biochemical functions such as autoinhibition^7^, subcellular localization, enzymatic activities or complex formation^8,9^. Understanding how inhibitors allosterically control such non-catalytic kinase functions is however key to understanding drug action and toxicity.

The KD of most kinases consists of conserved structural elements which constitute a conformational switch toggling between an active and inactive state. Human kinases are regulated via autoinhibitory domains (AIDs) which, in the absence of activating signals, keep kinases in an inactive state by binding to the KD and masking ATP binding site^10^, controlling the accessibility and conformation of the ATP-binding domain blocking both kinase activity and interactions with substrates^11,12^. Type I inhibitors stabilize the active state of the KD, also called the aspartate-phenylalanine-glycine (DFG)-in conformation. However, it remains unclear whether this active state also involves structural changes at the AID, since the probe might not bind the autoinhibited state where the AID also binds at the active site. This lack of data can be explained by the fact that structural changes in AID domains are difficult to characterize by classical structural methods as many AIDs are disordered^13,14^ or linked to the KD via a disordered linker^15^. Therefore, it is plausible that a large fraction of kinases may undergo larger domain-domain interaction and subsequent contextual changes in response to ATP-competitive inhibitor binding. Consequently, while the biochemical and cellular consequences resulting from inhibited substrate phosphorylation have been studied in great detail, the extent to which inhibitor-induced open conformations govern cellular drug phenotypes remains a major dark space in kinome pharmacology.

To bridge this gap and systematically discover such mechanisms, we combined structural and contextual proteomics and applied it to three disease associated kinases regulated by AIDs for which highly specific ATP competitive inhibitors were available: Calcium/calmodulin-dependent protein kinase kinase 2 (CAMKK2), Checkpoint kinase 1 (CHEK1), and Protein kinase C alpha (PRKCA). To resolve structural changes induced by ligand binding at high sequence coverage we employed limited proteolysis coupled with mass spectrometry (LiP-MS) on native affinity purified (AP) samples (AP-LiP-MS). We combined AP-LiP-MS with contextual proteomics, using both AP-MS^9^ and *in vivo* proximity labeling^16^ as complementary approaches^17^ to understand how inhibitor induced conformational shifts reshape the kinase’s interactome and biochemical environment. The combination of these methods provides detailed insights on structural and protein-protein interaction (PPI) changes in response to inhibitors without relying on prior knowledge and represents a generalizable and unbiased approach that guides the *de novo* discovery of mechanisms of action.

Using our approach we identified a structural mechanism shared across all three kinases where inhibitor binding displaces the AID inducing an active-like kinase conformation. This open but catalytically inhibited state triggers novel molecular and cellular phenotypes, such as complex remodelling, sequestration of proteins away from pathways or changes in protein localization. Our findings suggest that inhibitor-induced neomorphic functions may represent a prevalent feature among kinases with AIDs, establishing our multimodal proteomic strategy as a systematic roadmap to identify off-mechanism liabilities for the development of safer, more effective kinase-targeted therapeutics and probes.

## Results

### ATP-competitive kinase inhibition displaces the autoinhibitory domain to induce an open conformation

To investigate probe-induced conformational changes in flexible protein regions, we used LiP-MS^18^, a structural proteomics method that relies on the generation of conformation specific protein fragments by the protease proteinase K (PK), which preferentially cleaves accessible and flexible regions^19^. Perturbations that alter protein accessibility or flexibility lead to changes in PK cleavage patterns, resulting in differential peptide abundance. The sensitivity of detecting these structural changes critically depends on protein sequence coverage, which can limit the analysis of low-abundance proteins such as kinases in complex cell lysates^18^. While previous studies have used purified recombinant proteins to address this limitation^20,21^, this approach does not fully capture the native context of the protein since it lacks physiological PTMs and complex partners essential for correct structure and function. Therefore, we performed LiP-MS on native AP eluates (Fig. 1A). In this approach, target proteins are purified from human cells under native conditions and treated with an inhibitor, prior to the LiP-step with PK. Samples are then digested with trypsin and analyzed by liquid chromatography coupled mass spectrometry (LC-MS).

**Fig. 1.**
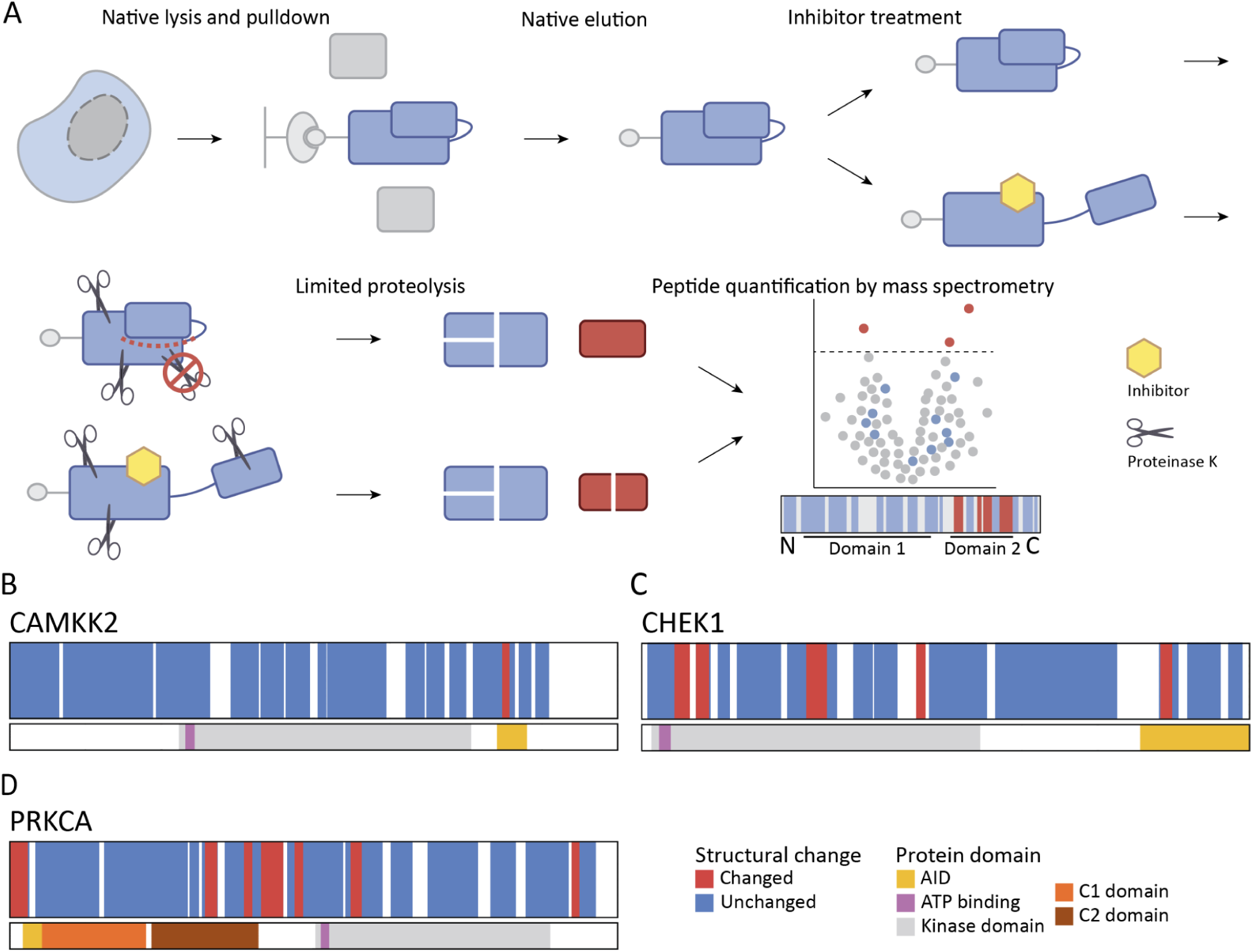
AP-LiP-MS analysis reveals ATP-competitive inhibition-induced structural changes in kinase AIDs. A) Overview of the AP-LiP-MS workflow. Proteins were lysed and eluted under native conditions before the corresponding inhibitor was added. Following limited proteolysis with PK, the conformation-specific protein fragments were further digested with trypsin under denaturing conditions. Finally, peptides were measured by LC-MS/MS and compared across conditions. B-D) Structural changes observed with AP-LiP-MS for CAMKK2 treated with SGC-CAMKK2-1, CHEK1 treated with rabusertib, and PRKCA treated with Gö 6983. In red are significantly changing peptides, in blue are the non-changing peptides. All AP-LiP-MS experiments were performed in triplicates and p-values were calculated using a moderated t-test followed by multiple testing corrections using the procedure of Benjamini-Hochberg.

We selected the serine/threonine-protein kinase DCLK1 to benchmark our approach, since X-ray crystal structures are available for both the DCLK1-IN-1-bound^22,23^ and unbound protein^24^ (Supplementary Fig. 1A). AP-LiP-MS profiling of DCLK1 showed that 25 of 29 peptides with significantly altered intensity were DCLK1 peptides, clearly indicating structural changes in DCLK1 upon inhibitor binding (Supplementary Fig. 1B, C, 2A). This observation shows that we detect the structural changes we anticipated from the X-ray structures.

**Fig. 2.**
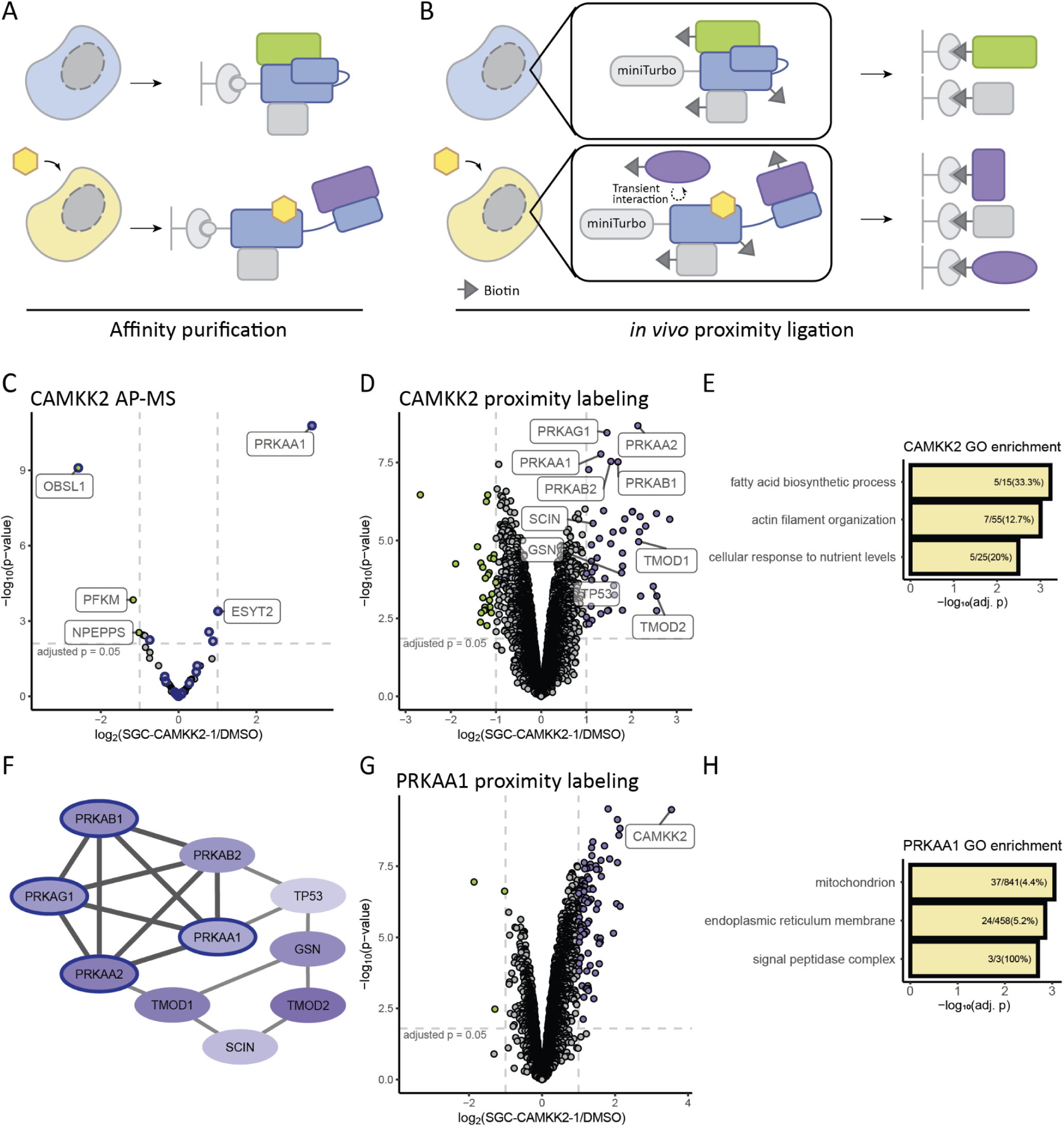
Binding of the inhibitor SGC-CAMKK2-1 changes CAMKK2 complex formation and proximity. A) Illustration of the AP-MS workflow. B) Illustration of the *in vivo* proximity labeling workflow. C) Volcano plot illustrating changes in CAMKK2 interactors upon inhibition with SGC-CAMKK2-1 measured by AP-MS. Known CAMKK2 interactors are highlighted with a blue border. P-values were calculated using a moderated t-test followed by multiple testing corrections using the procedure of Benjamini-Hochberg. D) Changes in the proximity of CAMKK2 upon inhibition with SGC-CAMKK2-1. P-values were calculated using a moderated t-test followed by multiple testing corrections using the procedure of Benjamini-Hochberg. E) GO enrichment analysis (biological process) of the significantly changing proteins in panel D. P-values were calculated using a Fisher’s exact test followed by multiple testing corrections using the procedure of Benjamini-Hochberg. F) PPI network showing the subunits of AMPK complexes and highly interconnected interactors that are significantly increased in the proximity of CAMKK2 upon treatment with SGC-CAMKK2-1 shown in panel D. The AMPK subunit interactions are highlighted with darker edges, and known interactors of CAMKK2 are outlined in blue. G) Reciprocal *in vivo* proximity labeling data showing changes in PRKAA1 proximity upon CAMKK2 inhibition. P-values were calculated using a moderated t-test followed by multiple testing corrections using the procedure of Benjamini-Hochberg. H) GO enrichment analysis (cellular component) of the significantly changing proteins in panel G. P-values were calculated using a Fisher’s exact test followed by multiple testing corrections using the procedure of Benjamini-Hochberg. AP-MS experiments were performed in triplicates, and *in vivo* proximity labeling experiments in quadruplicates.

To determine whether AID rearrangement upon inhibitor binding is a general phenomenon, we investigated kinases with diverse AIDs that vary in both structure and regulatory mechanism, and for which well-characterized ATP-competitive inhibitors are available. We therefore selected CAMKK2, CHEK1, and PRKCA as model kinases, representing distinct modes of autoinhibition and regulatory control. The AID of CAMKK2 is a small linear motif and binds the back of the KD^25^. Activation occurs through binding of calmodulin to the AID, which relieves the interaction between the AID and the KD. CHEK1 instead has a larger, structured AID. Its activation is triggered via phosphorylation from the upstream kinase Ataxia telangiectasia and Rad3 related (ATR), which relieves the interaction between the AID and the KD of CHEK1^26^. On the other hand, PRKCA has an N-terminal disordered pseudosubstrate motif that dissociates from the KD upon Ca^2+^ binding^27,28^. For each of these kinases, specific high-affinity ATP-competitive inhibitors have been developed: SGC-CAMKK2-1 for CAMKK2^29^, rabusertib for CHEK1^30^, and Gö 6983 for PRKCA^31^.

We applied AP-LiP-MS to CAMKK2, CHEK1 and PRKCA to investigate the structural consequences of inhibitor binding. For all three kinases we observed a structural change at the AID upon binding of the respective inhibitor (Fig. 1B-D, Supplementary Fig. 2B-G). CAMKK2 showed changes exclusively in its AID. The peptide indicating the structural change at the AID is half-tryptic, meaning that one end was generated by PK cleavage. The increased abundance of such peptides indicates increased PK susceptibility and thus increased accessibility of the AID. This is consistent with a transition to an open conformation. In addition to changes at the AID, CHEK1 and PRKCA also underwent structural alterations throughout the KD. Additionally, the PRKCA inhibitor Gö 6983 led to structural changes in the C2 domain, specifically at residues 260-269, which are part of a short sequence associated with autoinhibition^32^. We observed increased PK accessibility of both domains, potentially reflecting an allosteric disruption of the C2-KD interaction. Notably, all observed structural changes upon inhibitor treatment occurred in the same regions previously described to be involved in kinase activation. Specifically, the AID is known to dissociate from the KD upon activation for all investigated kinases^25,33–35^. Furthermore, PRKCA activation is also associated with larger rearrangements in its membrane-binding C2 domain^36^. These results suggest that kinases undergo conformational changes upon inhibitor binding in regions typically associated with their activation.

### Inhibition of CAMKK2 with SGC-CAMKK2-1 stabilizes interactions associated with CAMKK2 activation

An open kinase conformation induced by inhibitor binding could alter the accessibility of interaction surfaces hidden in the autoinhibited conformation^37^. We therefore applied AP-MS (Fig. 2A) and *in vivo* proximity labeling (Fig. 2B) to profile subsequent changes in kinase complexes and proximity, respectively.

We first measured changes in CAMKK2 complex formation. AP-MS identified 66 interactors of CAMKK2 across all treatments, 22 of which were previously listed in the BioGrid database^38^ (Supplementary Table 1). Five proteins changed in abundance in response to SGC-CAMKK2-1 (Fig. 2C, Supplementary Fig. 3A, B). 5’-AMP-activated protein kinase subunit alpha-1 (PRKAA1), the catalytic subunit of the trimeric AMPK complex and known substrate^39^ and interactor^40^ of CAMKK2, was the most strongly increased protein in CAMKK2 complexes in the presence of the inhibitor. Extended synaptotagmin-2 (ESYT2), another known CAMKK2 interactor, was also increased but to a lesser extent than PRKAA1. We also found three proteins whose interaction was reduced upon inhibitor binding. This includes a strong reduction of Obscurin-like protein 1 (OBSL1), a known CAMKK2 interaction albeit of unknown function and, with lower significance, the reduced abundance of metabolic enzyme ATP-dependent 6-phosphofructokinase (PFKM) and the Puromycin-sensitive aminopeptidase (NPEPPS).

To explore changes in the interaction landscape of CAMKK2 *in vivo*, we extended our analysis by proximity labeling using miniTurbo-CAMKK2 expressing cells (Fig. 2D, Supplementary Fig. 3C, D). We found 71 proteins with altered proximity to CAMKK2 in the presence of the inhibitor. Gene ontology (GO) enrichment analysis revealed that SGC-CAMKK2-1 caused changes in proximity to proteins related to actin filament organization and metabolic processes (Fig. 2E). Among these proteins were five subunits of the trimeric AMPK complex as well as known AMPK interactors, such as TP53 (Fig. 2F). The increased proximity to catalytic as well as regulatory AMPK subunits suggests complex formation between CAMKK2 and trimeric AMPK complexes upon SGC-CAMKK2-1 treatment. Interestingly, since the CAMKK2-PRKAA1 interaction has been previously observed only in the presence of Ca^2+^, calmodulin and ATP, this interaction likely requires an active CAMKK2 conformation^41^. Thus, our observation that CAMKK2 binds AMPK upon inhibition with SGC-CAMKK2-1 is consistent with the AP-LiP-MS data which suggested that the inhibitor stabilizes an open, active-like kinase conformation.

Reassuringly, SGC-CAMKK2-1 also induced a strong increase of endogenous CAMKK2 in the proximity of PRKAA1 following inhibitor treatment upon reciprocal *in vivo* proximity labeling using miniTurbo-PRKAA1 expressing cells (Fig. 2G, Supplementary Fig. 3E, F), confirming our previous findings. SGC-CAMKK2-1 treatment also caused a remodeling of PRKAA1 interactions beyond CAMKK2 as we observed a significant enrichment for proteins linked to mitochondria and endoplasmic reticulum based on a GO enrichment analysis (Fig. 2H). This suggests that CAMKK2 inhibitors may also change the biochemical environment of AMPK beyond stabilizing the CAMKK2-PRKAA1 interaction.

Taken together, we found that SGC-CAMKK2-1 causes structural changes and remodeling of CAMKK2-containing complexes, including a strong increase in CAMKK2-PRKAA1 complex formation. Consequently, probe treatment extensively altered the protein environment of PRKAA1. Our data thus show how an ATP-competitive kinase inhibitor can affect the interactors of both its target but also of associated signaling proteins.

### Sequestration of PRKAA1 by inhibitor-bound CAMKK2 inhibits PRKAA1 phosphorylation by other kinases

The inhibitor-induced CAMKK2-AMPK complex could be caused by the observed changes in CAMKK2 conformation but also result from decreased AMPK phosphorylation following catalytic inhibition of CAMKK2^42^. To disentangle the effects of catalytic inhibition and conformational change on the altered PPIs, we compared inhibitor-induced structural changes and complex formation between wildtype (WT) CAMKK2 and the following three mutants: the catalytically inactive D312A mutant^43^, the R311C mutant found in patients with bipolar disorder^44^, and the phosphomimetic mutation T483D, which is at the major autophosphorylation site of CAMKK2 associated with autonomous activity^45^.

First, we applied AP-LiP-MS to compare structural changes between wildtype and mutant CAMKK2 following inhibitor treatment. We found that all tested mutants displayed a structural change at or close to the AID similar to wildtype CAMKK2 in response to SGC-CAMKK2-1 (Fig. 3A, Supplementary Fig. 4A-C), demonstrating that all mutants can bind the compound and respond with a conformational change similar to WT CAMKK2. The structural change in the AID for the CAMKK2 T483D mutant suggests that it is also in a closed conformation in the absence of the inhibitor. This is consistent with previous reports showing that activity of the CAMKK2 isoform used in our study depends on calmodulin binding also when autonomously active^46^.

**Fig. 3.**
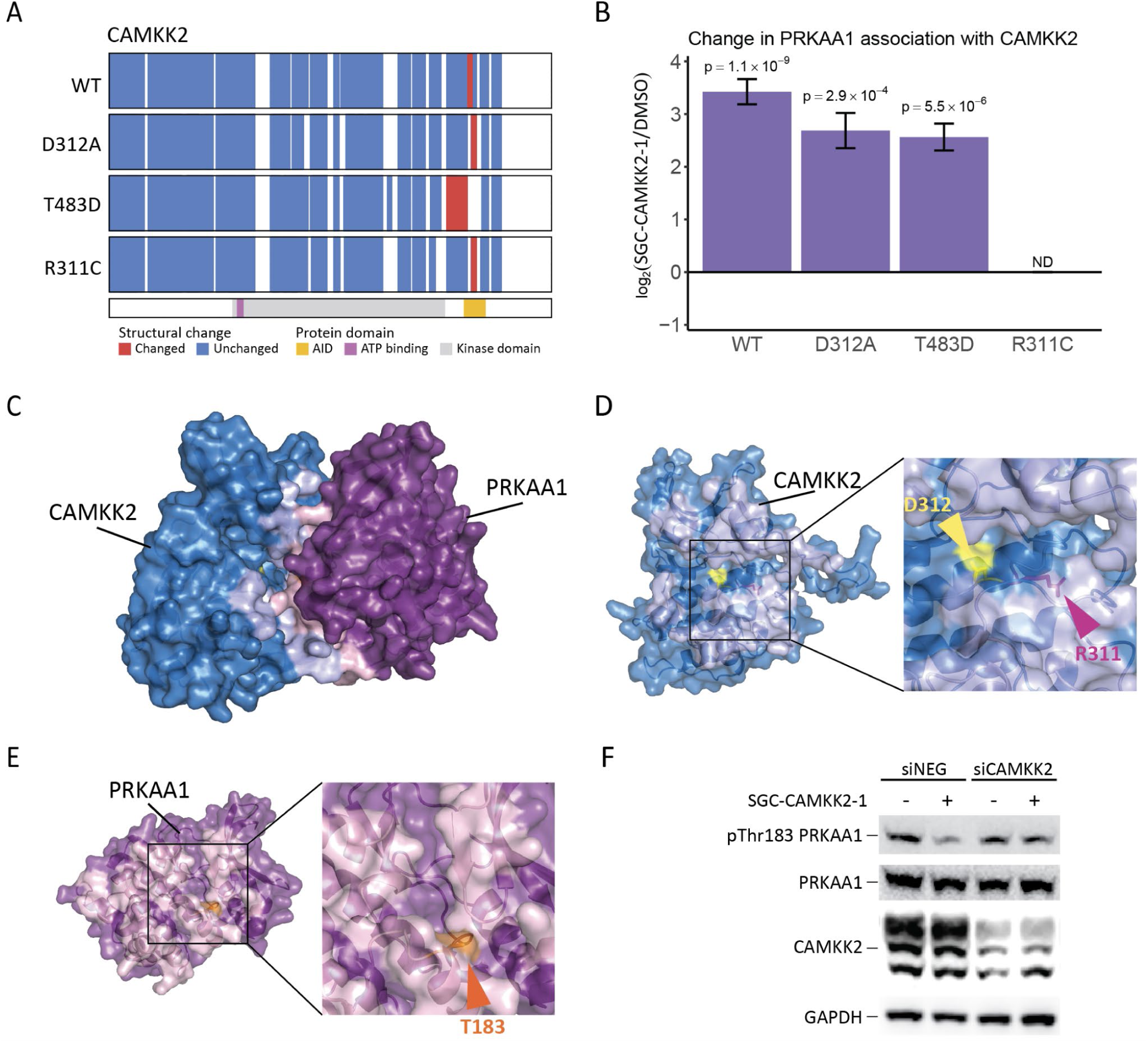
Inhibitor-induced CAMKK2-PRKAA1 complex formation can attenuate T183 phosphorylation of PRKAA1 by other kinases. A) Comparison of structural changes in CAMKK2 mutants upon SGC-CAMKK2-1 treatment as measured by AP-LiP-MS. B) Bar graph representing log_2_(fold change) in PRKAA1 association with CAMKK2 wildtype and indicated mutants following SGC-CAMKK2-1 treatment as measured by AP-MS. Error bars represent the standard error of the mean of the log_2_(fold change). C) Predicted structure of the CAMKK2-PRKAA1 complex. CAMKK2 is shown in blue, and PRKAA1 is shown in purple and the predicted interfaces are highlighted in light blue and pink, respectively. Only kinase domains were used in the prediction. D) An up-close view of R311 and D312 of CAMKK2 at the CAMKK2-PRKAA1 interface. R311 is shown in magenta, and D312 is shown in yellow. The predicted interface is highlighted in light blue. E) The predicted PRKAA1 interface in the CAMKK2-PRKAA1 complex. The residue T183 is shown in orange, the predicted interface highlighted in pink. F) Western blot of PRKAA1 phosphorylation in response to CAMKK2 inhibition or siRNA mediated CAMKK2 knockdown. HEK293 cells were treated with control siRNA (siNEG) or siCAMKK2 for 48 hours and vehicle control or SGC-CAMKK2-1 for 4 hours as indicated. Cells were then subjected to glucose starvation for 20 minutes to induce T183 phosphorylation on PRKAA1. AP-LiP-MS and AP-MS experiments were performed in triplicates. P-values were calculated using a moderated t-test followed by multiple testing corrections using the procedure of Benjamini-Hochberg.

We subsequently analyzed CAMKK2-PRKAA1 complex formation upon SGC-CAMKK2-1 treatment across the mutants (Fig. 3B, Supplementary Fig. 4D-F). Upon SGC-CAMKK2-1 treatment, both the catalytically inactive D312A and the autonomously active T483D mutant displayed an interaction pattern with PRKAA1 similar to wildtype CAMKK2, suggesting that these changes are not linked to CAMKK2 catalytic activity but rather relate to the observed structural change. Surprisingly, the disease-related CAMKK2 R311C mutation completely abolished the interaction with PRKAA1, irrespective of inhibitor treatment, which could hint at a novel disease mechanism via uncoupling the CAMKK2-AMPK signaling axis in patients with this genetic disorder.

To better understand potential implications of CAMKK2-PRKAA1 complex formation, we generated a structural prediction of the two KDs using AlphaFold3^47^ (Fig. 3C, Supplementary Fig. 4G, H). In the predicted PPI interface of CAMKK2, we observed that R311 is oriented towards the predicted interaction interface, while D312 is outside of the interaction interface (Fig. 3D). This suggests that the R311C mutation may negatively affect the integrity of the interaction interface needed for PRKAA1 binding and may explain why the SGC-CAMKK2-1-induced interaction with PRKAA1 is abolished for CAMKK2 R311C but not D312A.

We further investigated the predicted binding interface of PRKAA1 (Fig. 3E) and noticed that T183, shown in orange, is buried in the interface of the predicted CAMKK2-PRKAA1 complex. While this residue can be phosphorylated by CAMKK2 in response to increased intracellular Ca^2+^ ions, it is phosphorylated upon glucose starvation primarily by liver kinase B1 (LKB1)^48^. Since PRKAA1 is bound by CAMKK2 in response to SGC-CAMKK2-1, this site would become inaccessible to other upstream kinases. This led us to hypothesize that formation of the CAMKK2-PRKAA1 complex would inhibit PRKAA1 T183 phosphorylation by other kinases such as LKB1. We tested this hypothesis by analyzing T183 phosphorylation in HEK293 cells subjected to glucose starvation and siRNA-mediated depletion of CAMKK2 (Fig. 3F, Supplementary Fig. 5). We found that the glucose-induced T183 phosphorylation of PRKAA1 was indeed decreased after CAMKK2 inhibition with SGC-CAMKK2-1. Importantly, this decrease was rescued upon siRNA-mediated knockdown of CAMKK2, supporting a model where SGC-CAMKK2-1 induced formation of CAMKK2-PRKAA1 complexes could block the T183 phosphorylation and activation of PRKAA1 by other kinases.

We showed that inhibitor-induced changes in protein interactions cannot be simply explained by inhibition of catalytic activity, and that SGC-CAMKK2-1 can regulate a non-catalytic scaffolding function of CAMKK2 via conformational changes. The disease-associated mutant R311C fully inhibited CAMKK2-PRKAA1 complex formation, representing first evidence for a potential molecular mechanism associated with bipolar disorder linked to this mutation and emphasizing the importance of genetic context for personalized drug response. Finally, we showed that SGC-CAMKK2-1 induces CAMKK2-AMPK complex formation, which may represent an additional mechanism to hinder PRKAA1 phosphorylation by other kinases. This finding underscores the importance of exploring inhibitor-induced PPI changes to broaden our understanding of multilayered target engagement mechanisms.

### Inhibition of CHEK1 with rabusertib alters interactions with DNA-damage response proteins

AP-LiP-MS revealed significant structural alterations in the KD and the AID of CHEK1 upon binding of the inhibitor rabusertib. To understand how these changes may remodel the interactome of CHEK1, we performed AP-MS and *in vivo* proximity labeling experiments. AP-MS identified 51 interactors of CHEK1, of which 39 have been reported previously^38^ (Supplementary Table 2). Rabusertib affected CHEK1 interactions with 18 proteins (Fig. 4A, Supplementary Fig. 6A) that upon GO enrichment analysis show clear enrichment for terms related to known CHEK1 functions in the DNA-damage response (Fig. 4B). This includes increased interaction with 14-3-3 proteins zeta (YWHAZ) and theta (YWHAQ), which has also been observed upon DNA damage-induced CHEK1 activation^49^. Inhibitor treatment in addition stabilized the interaction with Ubiquitin carboxyl-terminal hydrolase 7 (USP7), a deubiquitylating (DUB) enzyme known to directly interact with CHEK1^50^. USP7 has increased DUB activity towards CHEK1 upon DNA damage^51^. Moreover, rabusertib increased the abundance of CHEK1 complexes containing proliferating cell nuclear antigen (PCNA) and the DNA replication licensing factors MCM2, MCM3, MCM4, MCM5 and MCM6, which are subunits of the MCM complex. Both these interactions are required for CHEK1 activation^52,53^. These proteins are part of a highly interconnected PPI network (Fig. 4C). The only protein we found to dissociate from CHEK1 upon rabusertib treatment was the mitochondrial protein CLPB.

**Fig. 4.**
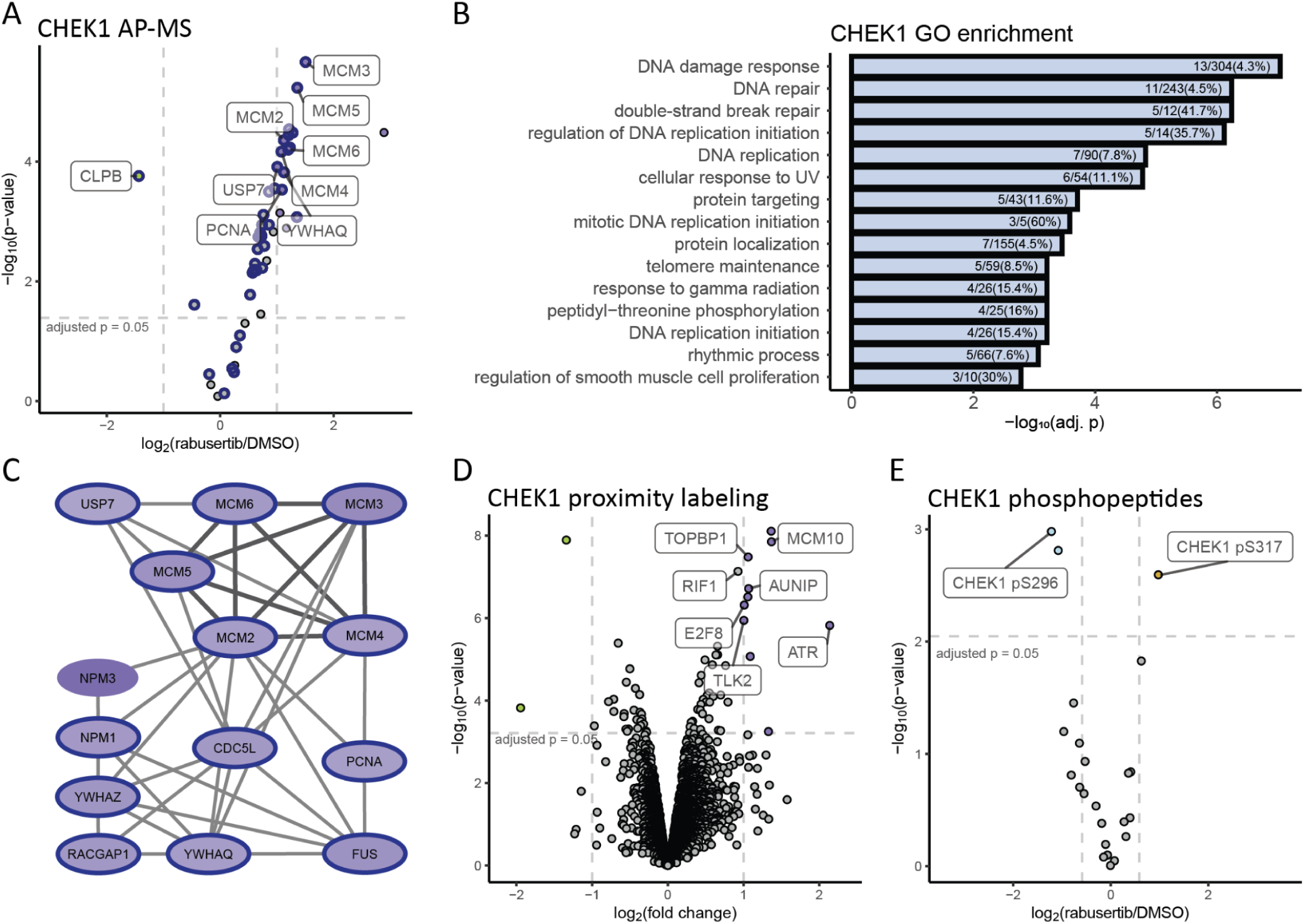
CHEK1 inhibition with rabusertib alters its interaction with multiple DNA-damage response proteins. A) Volcano plot illustrating changes in abundance of CHEK1 interactors in response to rabusertib identified by AP-MS. Known CHEK1 interactors are highlighted with a blue border. P-values were calculated using a moderated t-test followed by multiple testing corrections using the procedure of Benjamini-Hochberg. B) GO enrichment analysis (biological process) of all interactors of CHEK1 identified by AP-MS. P-values were calculated using a Fisher’s exact test followed by multiple testing corrections using the procedure of Benjamini-Hochberg. C) A highly interconnected subnetwork of the proteins associating with CHEK1 upon rabusertib treatment measured in panel A. The MCM complex is highlighted with darker edges, and known interactors of CHEK1 are highlighted with a blue border. D) Changes in the proximity of CHEK1 upon inhibition with rabusertib. P-values were calculated using a moderated t-test followed by multiple testing corrections using the procedure of Benjamini-Hochberg. E) Changes in phosphorylation upon CHEK1 inhibition with rabusertib extracted from AP-MS results. P-values were calculated using a moderated t-test followed by multiple testing corrections using the procedure of Benjamini-Hochberg. AP-MS experiments were performed in triplicates, and *in vivo* proximity labeling experiments in quadruplicates.

*In vivo* proximity labeling with miniTurbo-CHEK1 showed that rabusertib induced the proximity to 12 proteins with known functional links to CHEK1 (Fig. 4D, Supplementary Fig. 6B, C) including Aurora kinase A- and ninein-interacting protein (AUNIP), transcription factor E2F8, Rap1-interacting factor 1 (RIF1), MCM10, the kinases tousled-like 2 (TLK2) and ATR, and the scaffolding protein DNA topoisomerase 2-binding protein 1 (TOPBP1). ATR and TOPBP1 act upstream to activate CHEK1 by phosphorylation on multiple sites including S317 upon DNA damage^54^.

We therefore analyzed the abundance of phosphopeptides in purified CHEK1 complexes, and observed increased phosphorylation of CHEK1 on the known ATR phosphorylation site S317 upon rabusertib treatment (Fig. 4E, Supplementary Fig. 6D). This paradoxical activating phosphorylation in response to CHEK1 inhibition has been noticed in the past and might be due to increased spontaneous DNA damage as observed previously for other CHEK1 inhibitors^55,56^ (Supplementary Fig. 6E, 7). We also observed decreased CHEK1 autophosphorylation at S296 (Fig. 4E) in rabusertib-treated cells as expected upon CHEK1 catalytic inhibition^57^. These phosphorylation changes might explain both the enhanced 14-3-3 protein binding to CHEK1, which depends on phosphorylation by ATR^49^, and the increase in binding to chromatin and associated proteins, as CHEK1 dissociation from these depends on S296 autophosphorylation^58^.

We found that CHEK1 inhibition changes CHEK1 complex formation with many DNA-damage response proteins. These changes were consistent with decreased CHEK1 activity, resulting in lower levels of autophosphorylation and increased DNA damage and thus increased ATR activity towards CHEK1. We show that such mechanisms can also shape the interaction landscape of an inhibited kinase.

### CHEK1 inhibition destabilizes complexes with the mitochondrial protein CLPB and causes mitochondrial fragmentation

Whereas the functional relevance of most of the observed interactors has been studied, the role of the CHEK1-CLPB interaction change remains unknown. Several studies using different approaches have identified CLPB as an interaction partner of CHEK1^9,59,60^ (Fig. 5A), which strongly suggests the formation of mitochondrial CHEK1 complexes of yet unknown function. We were therefore prompted to study the rabusertib-induced CHEK1-CLPB dissociation in more detail. To test whether the dissociation depends on the catalytic activity of CHEK1, we compared dissociation in WT CHEK1, the catalytically inactive D130A mutant and the constitutively active L449R mutant, which is in an open conformation that is constitutively phosphorylated by ATR^33,61^. Rabusertib treatment significantly decreased CLPB in both WT and D130A CHEK1 complexes, whereas CHEK1 L449R-CLPB binding was not affected (Fig. 5B, Supplementary Fig. 8A, B). This suggests that the interaction with CLPB does not depend on catalytic inhibition of CHEK1.

**Fig. 5.**
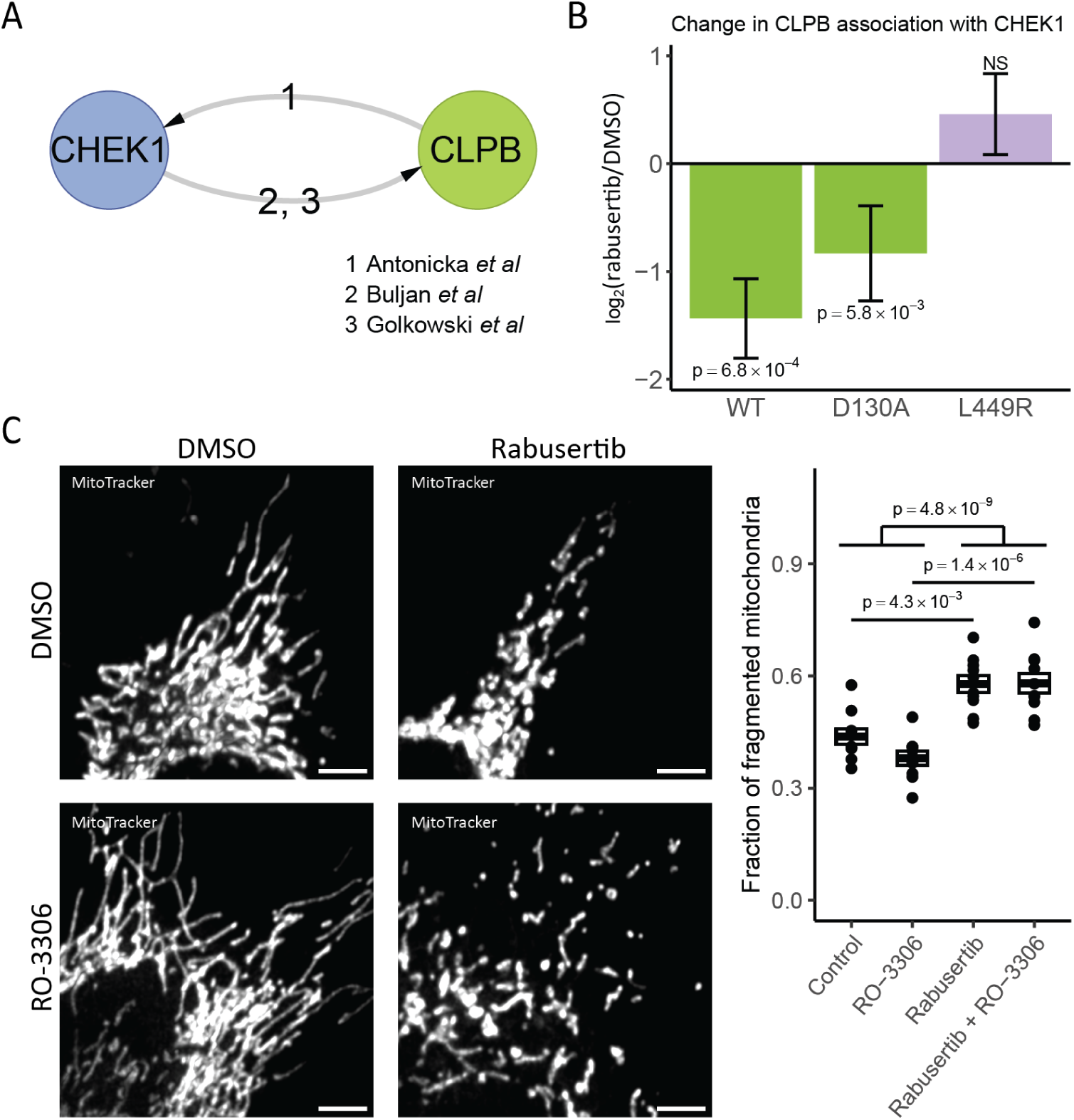
Rabusertib disrupts CHEK1-CLPB complex formation and causes mitochondrial fragmentation. A) Previous studies detecting CHEK1-CLPB complex. B) Bar graph representing log_2_(fold change) in CLPB association with CHEK1 WT and indicated mutants upon rabusertib treatment. Error bars represent the standard error of the mean of the log_2_(fold change). P-values were calculated using a moderated t-test followed by multiple testing corrections using the procedure of Benjamini-Hochberg. Experiments were performed in triplicates. C) Super-resolution microscopy of mitochondria in HeLa cells treated with rabusertib in the presence and or absence of RO-3306. Images were segmented and mitochondria were classified based on their morphology as either fragmented or elongated. Within each image, the area of fragmented mitochondria was normalized by the total area of detected mitochondria. P-values were calculated by ANOVA followed by Tukey HSD. For each condition, 10 images taken across 3 replicates were quantified. The scale bars represent 5 μm.

Given that genetic deletion of CLPB results in mitochondrial fragmentation^62^, we next analyzed mitochondrial morphology in fixed HeLa cells in the presence or absence of rabusertib using super-resolution microscopy (Fig. 5C). We quantified mitochondria in fragmented and elongated states and identified mitochondrial areas consistent with previous analyses of human mitochondria^63,64^ (Supplementary Fig. 8C). Rabusertib treatment caused a significant increase in mitochondrial fragmentation (p = 4.3 × 10^-3^, ANOVA followed by Tukey HSD). CHEK1 inactivation can lead to premature and sustained CDK1 activation^65^, which induces mitochondrial fragmentation via phosphorylation of the GTPase Dynamin-1-like protein (DRP1) controlling mitochondrial fission^66,67^. To exclude such a mechanism, we analyzed the mitochondrial network in response to rabusertib in combination with the CDK1 inhibitor RO-3306. We confirmed the significant increase in mitochondrial fragmentation even when CDK1 is inhibited by RO-3306 (p = 1.4 × 10^-6^, ANOVA followed by Tukey HSD), which excludes mitochondrial fragmentation via activation of the CDK1-DRP1 axis.

We had noticed that CHEK1 inhibition by rabusertib can cause DNA damage similar to etoposide treatment as measured by increased histone H2AX phosphorylation (Supplementary Fig. 6E, 7). We therefore asked if mitochondrial fragmentation could also be detected upon etoposide-mediated DNA damage. We observed a slight but non-significant increase in mitochondrial fragmentation upon etoposide treatment (p = 0.151, two-sided t-test; Supplementary Fig. 8D). This indicates that DNA damage alone cannot explain mitochondrial fragmentation by rabusertib, suggesting the involvement of an additional mechanism. In this regard, rabusertib-induced dissociation of mitochondrial CHEK1-CLPB complexes represents an intriguing mechanism, though further testing is needed before a causal relationship can be established.

Taken together, we observed that CHEK1 inhibition is associated with changes in PPIs, including dissociation of the known mitochondrial interactor CLPB. We showed that the interaction between CHEK1 and CLPB does not depend on catalytic activity of CHEK1. Based on this finding, we investigated mitochondrial phenotypes upon CHEK1 inhibition. We found that rabusertib induced a significant increase in mitochondrial fragmentation upon CHEK1 inhibition by a mechanism that is independent of CDK1 activation or induction of DNA damage. This illustrates how non-catalytic probe mechanisms can induce novel phenotypes.

### Inhibitor binding induces relocalization of PRKCA to cellular junctions

We found that upon Gö 6983 treatment, PRKCA undergoes structural rearrangements in the KD and AID, but also in other domains, such as the membrane-associating C2 domain. We aimed to profile related interactome changes of PRKCA using AP-MS. Following purification from native lysates obtained from treated cells, we surprisingly noticed a four-fold decrease in PRKCA abundance upon Gö 6983 treatment (Supplementary Fig. 9A, B), limiting the sensitivity of the AP-MS approach. In contrast, we observed no such decrease in PRKCA abundance in the *in vivo* proximity labeling experiment, which applies harsh denaturing lysis conditions, suggesting that Gö 6983 treatment results in PRKCA relocalization to a cell compartment that is insoluble under native lysis conditions.

To characterize the relocalization of inhibitor-bound PRKCA, we focused our analysis on *in vivo* proximity labeling. This approach clearly demonstrated a Gö 6983 induced shift in PRKCA proximity towards membrane associated proteins linked to cell adhesion and cellular junctions (Fig. 6A-C, Supplementary Fig. 9C, D). This included the focal adhesion protein and known PRKCA interactor Integrin beta-1 (ITGB1)^68^, but also proteins from tight junctions (OCLN, MARVELD2, CXADR, FMN2, EPB41L4B), adherens junctions (NECTIN2, SNAP23), desmosomes (DSG2) and other junction proteins (ADM4, KIRREL1, MPZL1, TENM3), suggesting that PRKCA relocalizes to very specific membrane structures that control cell-cell contacts and intercellular signaling upon Gö 6983 binding.

**Fig. 6.**
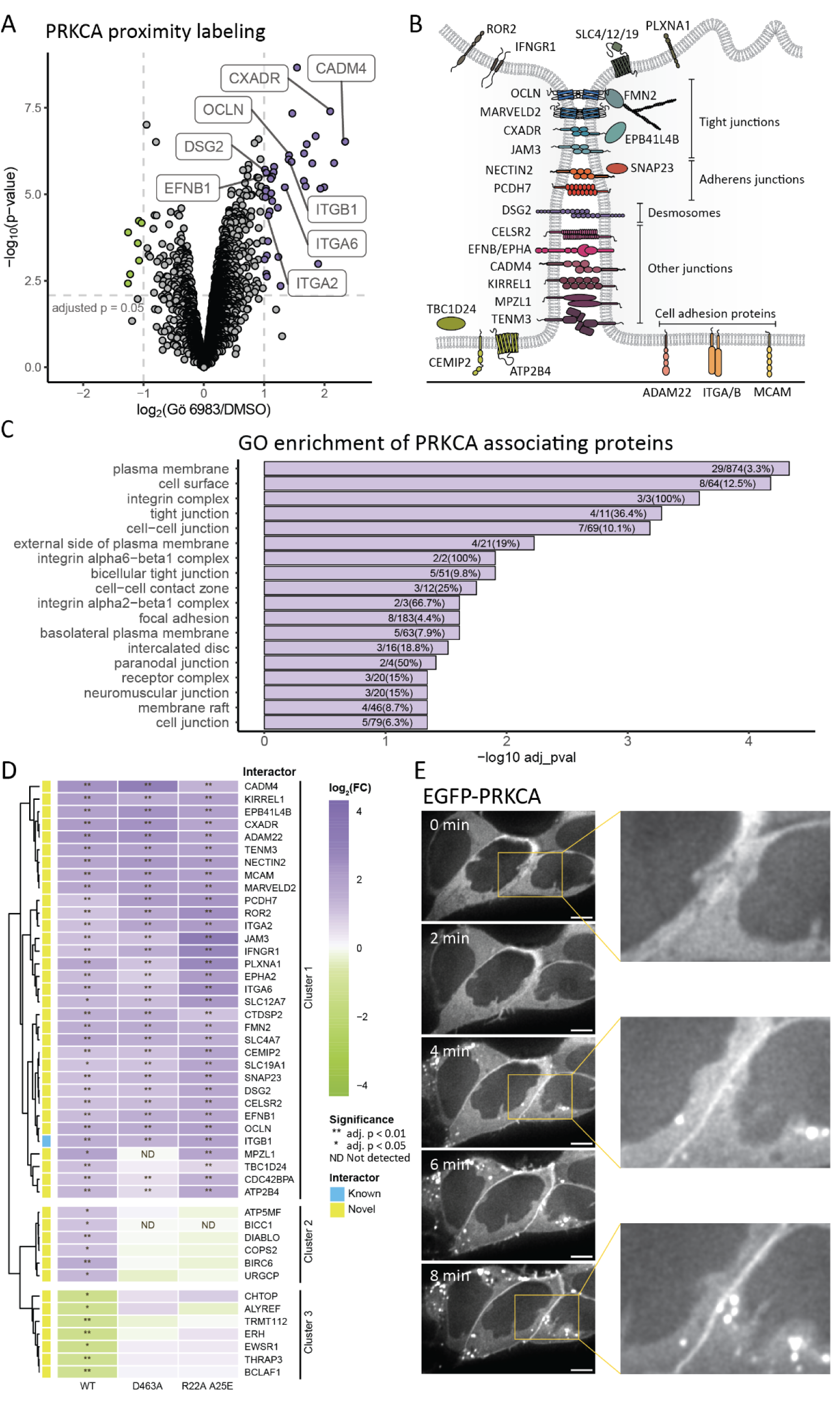
PRKCA changes localization to cellular junctions upon its inhibition with Gö 6983. A) Changes in the proximity of PRKCA upon inhibition with Gö 6983. P-values were calculated using a moderated t-test followed by multiple testing corrections using the procedure of Benjamini-Hochberg. B) Illustration of the proteins annotated with GO terms ‘plasma membrane’ or ‘cell adhesion’ found increased in proximity to PRKCA upon Gö 6983 treatment. C) GO enrichment analysis (cellular component) of proteins associating with PRKCA upon Gö 6983 treatment. P-values were calculated using a Fisher’s exact test followed by multiple testing corrections using the procedure of Benjamini-Hochberg. D) Comparison of proximity changes upon Gö 6983 treatment in PRKCA WT, the catalytically inactive D463A mutant and the active R22A/A25E double mutant. P-values were calculated using a moderated t-test followed by multiple testing corrections using the procedure of Benjamini-Hochberg. E) Super-resolution live-cell microscopy of HEK293 cells expressing EGFP-PRKCA upon Gö 6983 treatment. The scale bars represent 5 μm.

Under physiological conditions, the affinity of PRKCA for membrane lipids is regulated by Ca^2+^ binding^69^. We therefore asked if the observed relocalization to membrane structures could be explained via changes in intracellular Ca^2+^ concentrations induced by Gö 6983. We however did not observe any changes in cellular Ca^2+^ concentration upon Gö 6983 treatment using a Fluo-4 calcium imaging assay (Supplementary Fig. 9E). We therefore excluded the possibility that Gö 6983 alters the biochemical context of PRKCA via changes in Ca^2+^ concentration, and proposed a model where Gö 6983 binding to PRKCA directly affects the affinity of the kinase for membranes or membrane proteins.

To explore whether catalytic activity is needed for the observed proximity changes, we compared Gö 6983-induced proximity patterns between WT PRKCA, the catalytically inactive PRKCA D463A mutant and the constitutively active R22A/A25E double mutant^70^. When inhibited, we found similar changes in the proximity of PRKCA WT and both mutants towards the cellular junction proteins (Fig. 6D, cluster 1, Supplementary Fig. 9F, G), suggesting that these changes are independent of catalytic inhibition. Other proximity changes were affected in the mutants compared to the WT (Fig. 6D, cluster 2 and 3), suggesting that these changes may depend on the catalytic activity or the conformation of the AID. We next compared our biochemical findings to subcellular localization patterns using live-cell imaging of cells expressing EGFP-PRKCA. Indeed, consistent with our proximity data, we observed Gö 6983-induced relocalization of PRKCA towards the cell periphery, especially at cell-cell contact sites, within 8 minutes (Fig. 6E). In addition to the increased interaction with the plasma membrane, we also noticed localization towards unknown cytoplasmic foci. Since we saw no enrichment for proteins associated with intracellular organelles, these foci might represent PRKCA aggregates or endosomes containing cell-junction proteins^71^.

In summary, we observed that Gö 6983 caused a rapid relocalization of PRKCA towards the plasma membrane, specifically towards complexes involved in cell adhesion and cellular junctions. These changes were observed also in the catalytically inactive and constitutively active mutants upon Gö 6983 treatment, suggesting that these changes are not related to the inhibition of PRKCA kinase activity. PRKCA is known to associate with the plasma membrane through direct contact with lipids upon activation via its C2 domain^72^. Since our AP-LiP-MS data suggest that Gö 6983 binding can trigger structural changes in the C2 domain, we propose that these structural changes expose the C2 domain for subsequent recruitment of PRKCA to specific membrane structures.

## Discussion

ATP competitive kinase inhibitors are primarily designed to block the catalytic activity of their target, but their effect on kinase conformation and non-catalytic functions remains poorly explored. X-ray crystallography studies revealed that type I and II inhibitors can stabilize either an active DFG-in or inactive DFG-out conformation of the KD, respectively^73^. The DFG-in and DFG-out classification, although initially useful, turned out to be insufficient to fully explain drug action and cellular phenotypes. Our study suggests that the ATP-binding site is a major organizing center of kinase conformation and interaction, regulating contacts with AIDs, other regulatory domains and even other proteins. ATP-competitive inhibition disrupts these domain-domain interactions, resulting in active-like kinase configurations that drive molecular events including regulation of non-catalytic functions like complex formation or relocalization.

Using AP-LiP-MS, we detected structural changes distal from the orthosteric binding sites for all kinases analyzed, suggesting that inhibitor-induced allostery is a more pervasive feature of type I inhibitor binding. These changes are best explained by dissociation of the AID from the KD, a mechanism that mimics the conformational unlocking observed during physiological kinase activation. The observation that the AID is more solvent-exposed upon inhibitor binding and thus in the open conformation, together with previous reports that these inhibitors stabilize the active, DFG-in conformation of the KD^29,74,75^, indicates that the inhibitor-bound conformations closely resemble the active kinase structure. This is corroborated by our finding that the probe-bound conformation facilitates the formation of kinase complexes typically found only upon activation of CAMKK2, CHEK1 and PRKCA. Contextual proteomics revealed important new insights into neomorphic drug effects for all three models tested. In our first model, the CAMKK2 inhibitor SGC-CAMKK2-1, besides blocking CAMKK2 kinase activity, induced stable complex formation with PRKAA1. Our data and structural modeling suggest that SGC-CAMKK2-1 locks CAMKK2 in a conformation that sequesters PRKAA1, shielding its Thr183 from phosphorylation by upstream kinases like LKB1. Therefore, SGC-CAMKK2-1 binding may result in a pan-AMPK inhibition which shuts down both the calcium branch (via direct catalytic inhibition) and the energy stress arm (via sequestration). Such a dual AMPK inhibitor role may offer novel means to modulate AMPK-dependent energy signaling in tissues expressing CAMKK2 such as the central nervous system.

In our second model, CHEK1 inhibitor rabusertib disrupted CHEK1 complex formation with CLPB, a known CHEK1 interactor controlling mitochondrial homeostasis and stress responses^9,59,60^. While the precise function of this CHEK1-CLPB interaction remains to be fully resolved, it may regulate mitochondrial integrity. CLPB is known to maintain the solubility of the anti-apoptotic regulator HAX1^76^—which interacts with both CHEK1^9,60,77^ and CLPB^59,77,78^—allowing HAX1 to inhibit Caspase-9^79^. Furthermore, loss of CLPB has been linked to mitochondrial fragmentation^62^. Strikingly, we observed that rabusertib disrupts the CHEK1-CLPB interaction and causes mitochondrial fragmentation, even when CDK1 is inhibited. This finding rules out the canonical mechanism where CHEK1 inhibition drives fragmentation via promoting CDK1-mediated DRP1 activation^66,67^. Instead it suggests a mechanism independent of CHEK1 catalytic activity, potentially involving OPA1, an established client of CLPB^78^. In the absence of CLPB, OPA1 aggregates and levels are greatly reduced resulting in mitochondrial fragmentation^80^. While a direct functional link remains to be characterized, these findings suggest that CHEK1 may act as a regulator for the CLPB-mediated proteostasis machinery, effectively coupling the DNA damage response to the maintenance of mitochondrial structural integrity.

Finally, the PRKCA example illustrates how inhibitor-modulated domain-domain interactions can redirect PRKCA to specific subcellular compartments. As with the other kinases, we detected structural changes in the AID of PRKCA. Additionally, we also identified structural changes in a segment required for PRKCA autoinhibition within the C2 domain^32^, suggesting that multiple autoinhibitory domain-domain interactions are disrupted by Gö 6983 binding. The C2 domain is also required for membrane recruitment following PRKCA activation^69,81^. Consistent with these structural alterations, contextual proteomics indeed showed that inhibitor binding strongly induced the proximity of PRKCA to specific membrane proteins at cellular junctions, a phenotype we validated using live-cell imaging. Membrane recruitment under physiological conditions occurs upon PRKCA activation in response to Ca^2+^ and requires both the C1 and the C2 domain. Whereas C1 domain binds DAG^82^, it has been proposed that the C2 domain can bind phosphatidylserine (PS) upon Ca^2+^ signaling and phosphatidylinositol 4,5-bisphosphate (PIP2) via a lysine cluster^83,84^. Given that PIP2 is concentrated at the apical surface and tight junctions^85^ it is plausible that PRKCA recruitment to cell junction complexes may result from inhibitor induced opening of the C2 domain, which exposes the C2 lysine cluster for subsequent PIP2 binding at cellular junctions. Besides PIP2 tethering, it is also possible that PRKCA may be stabilized at the membrane via direct PPIs, for example via binding ITGB1^68^. Although the downstream effects of relocalizing inhibited PRKCA to cellular junctions are not yet fully understood, this potential for disrupting junctional architecture and intercellular signaling necessitates a broader evaluation of PRKCA inhibitors beyond their primary catalytic targets before their clinical application.

All three examples illustrate the breadth of catalysis-independent effects that can be induced upon type I inhibitor binding. They also suggest that inhibitor-induced non-catalytic gain-of-function is a more frequent phenomenon than currently anticipated. The majority of the 94 FDA approved kinase inhibitors are type I inhibitors^86^, yet we have reliable information on non-catalytic effects for only a few of these drugs. For instance, the FDA approved JAK2 inhibitor ruxolitinib prevents the interaction with tyrosine phosphatases by altering the conformation of the KD resulting in paradoxical hyperphosphorylated JAK2^87^. This primes JAK2 and causes a sudden increase in JAK2 activity after inhibitor dissociation, explaining major side effects in patients after drug withdrawal^88^. Another prominent example is the type I BRAF inhibitor GDC-0879, which paradoxically causes activation of downstream mitogen-activated protein kinase (MAPK) signaling and cell proliferation via an allosteric mechanism promoting dimerization between BRAF and CRAF^89,90^ even though the inhibitor was designed to inhibit this pathway. These paradoxical inhibitor responses were largely discovered serendipitously at late stages or post-approval.

The limited number of currently reported allosteric inhibitor responses reflects the limitations and inherently low throughput of traditional structural methods. Previous analyses often used highly purified, and usually truncated, recombinant proteins which lack key interaction partners, PTMs and domains. Multimodal proteomic target engagement analysis as presented here represents an important step to overcome previous limitations by enabling high throughput detection and characterization of allosteric inhibitor effects across the kinase family and beyond. Systematic characterization of such mechanisms during early drug development could help to reduce the high failure rate and could guide the development of safer and more effective therapeutics.

We therefore advocate for the systematic multiproteomic characterization of such off-mechanism effects during inhibitor development. Recent advances in LC-MS instrumentation now provides the scalability needed for systematic target engagement profiling^91^. Crucially, the combination of structural and contextual proteomics is not restricted to kinases and can be applied to a broad range of targets to develop new classes of inhibitors targeting also proteins with no known catalytic activity. By systematically mapping inhibitor modulated conformational and interactome landscapes, we can better understand drug phenotypes and reduce the failure rate of future therapeutics by detecting neomorphic liabilities early during drug development.

## Methods

### Generation of expression constructs

We generated doxycycline-inducible expression vectors of N-terminal Strep-HA-tagged, miniTurbo-tagged or EGFP-tagged bait proteins using the destination vectors pcDNA5-FRT-TO-SH-GW^92^, pcDNA5-pDEST-FRT-MiniTurbo-Flag-N-term, or pcDNA5-pDEST-FRT-EGFP-Flag-N-term, respectively. Human ORFs were provided as pDONR223 vectors selected from addgene, Gateway compatible human orfeome collections horfeome v5.1, horfeome v8.1 and ORFeome Collaboration Clones^93^ (OpenBiosystems, Horizon Discovery) or from a kinome orf collection^94^ (Supplementary Table 3). Mutations were introduced using a QuickChange II Site-Directed Mutagenesis Kit (Agilent) with primers listed in Supplementary Table 4. The LR reaction was done as described previously^95^. In brief, 150 ng of the destination vector and 300ng of the pDONR223 vector were mixed and diluted with TE buffer (pH 8.0) to a total volume of 4 µL. LR reaction was performed with 1 µL of LR Clonase II enzyme mix (Thermo Fisher Scientific) at 25°C for 1 h and transformed into OmniMAX competent *E. coli* cells (Thermo Fisher Scientific). The transformation mixture was plated onto LB plates containing 100 µg/ml ampicillin and incubated at 37°C o/n and plasmid DNA was isolated from individual colonies by using a QIAprep Spin Miniprep kit (Qiagen) according to the manufacturer’s instructions and confirmed by sequencing.

### Generation of stable inducible cell lines

The stable inducible cell lines were generated as described previously^95^ with the following alterations: 3 x 10^5^ T-Rex-HEK293 Flp-In cells (R78007, Invitrogen) were seeded onto six-well plates in DMEM supplemented with 10% fetal bovine serum (FBS) and 1% penicillin-streptomycin (P/S) and incubated at 37°C and 5% CO_2_ o/n. The cells were transfected with the following mixture: 100 ng of the respective expression vector, 900 ng of pOG44 (Invitrogen), 5.7 µl x-tremeGENE 9 DNA transfection reagent (Roche) and 94.3 µl DMEM. The mixture was incubated for 15 minutes at room temperature and added to the cells. The next day, new DMEM supplemented with 10% fetal bovine serum (FBS) and 1% penicillin-streptomycin (P/S) was added. The following day the cells were split into a T150 flask with a total volume of 32 ml. The next day hygromycin B was added at a final concentration of 100 µg/ml and hygromycin resistant recombinant cell pools were selected. Each new cell line was mycoplasma tested before use in experiments.

### Cultivation of cells for AP-LiP-MS and AP-MS experiments

HEK293 cells inducibly expressing the strep-HA-tagged bait proteins or EGFP (for AP-MS only) were seeded at a density of 10^7^ in 25 ml DMEM supplemented with 10% FCS (BioConcept) and 1% P/S in a 15 cm dish. The cells were incubated at 37°C and 5% CO_2_ o/n. Protein expression was induced with 1.3 μg/ml doxycycline for 24 hours before treatment with 1 μM inhibitor (SGC-CAMKK2-1: Sigma; rabusertib: Med Chem Express; Gö 6983: Med Chem Express) or 0.01% DMSO for 4 hours. Per sample, cells from two 15 cm dishes were harvested by scraping in cold PBS. Each sample was washed in PBS and flash frozen in liquid nitrogen.

### Cultivation of cells for proximity labeling experiments

HEK293 cells inducibly expressing the miniTurbo-tagged bait proteins or EGFP were seeded at a density of 10^7^ in 25 ml DMEM supplemented with 10% dialyzed FCS (BioConcept) and 1% P/S in a 15 cm dish. The cells were incubated at 37°C and 5% CO_2_ o/n. Protein expression was induced with 1.3 μg/ml doxycycline for 24 hours before they were treated with 1 μM inhibitor or 0.01% DMSO for 4 hours. An hour before harvesting, cells were treated with 50 μM biotin. Cells were harvested by dissociation in cold PBS with 1 mM EDTA. Each sample was washed in PBS and flash frozen in liquid nitrogen.

### AP-LiP-MS

Pellets were lysed by resolubilizing in 2 ml HNN buffer (50 mM HEPES pH 8.0, 150 mM NaCl, 50 mM NaF) supplemented with 0.5% IGEPAL, 400 nM Na_3_VO_4_, 1 mM PMSF and 0.2% Protease Inhibitor Cocktail (Sigma). The lysate was incubated on ice for 10 minutes and centrifuged at 18000x g for 20 minutes at 4°C. The supernatant was added to 80 μl StrepTactin Sepharose slurry (IBA Lifesciences) and incubated on a rotating wheel for 60 minutes at 4°C. The beads were loaded onto a 1 μm glass filter column and washed 3 times with 1 ml HNN buffer supplemented with 0.5% IGEPAL and 400 nM Na_3_VO_4_, 3 times with 1 ml HNN buffer, and 3 times with LiP buffer (1 mM MgCl_2_, 150 mM KCl and 100 mM HEPES pH 7.5). Samples were eluted twice by incubation with 30 μl of 0.5 mM biotin in LiP buffer for 15 minutes. The eluate was pooled and the total protein concentration was determined using a BCA protein assay kit (Pierce) according to the manufacturer’s protocol. Measured protein concentrations were between 0.06 μg/μl and 0.1 μg/μl. For each sample, 50 μl eluate was used. Samples were then processed as previously described^96^. Briefly, samples were treated with 10 μM inhibitor or 2% DMSO in triplicates and incubated at 25°C for 5 minutes. Proteinase K (Sigma) at an enzyme:substrate ratio of 1:100 was added for 5 minutes. Proteolysis was stopped by incubation at 99°C for 5 minutes. Samples were then diluted with 10% sodium deoxycholate (DOC) for a final concentration of 5% DOC. Samples were reduced with 5 mM TCEP for 40 minutes at 37°C and 200 rpm, and alkylated with 40 mM IAA for 30 minutes at 30°C in the dark, at 200 rpm. Samples were diluted with 100 mM ammonium bicarbonate to a DOC concentration of 1%. To each sample, 1 μg Lys-C and 1 μg trypsin was added and samples were incubated o/n at 37°C and 200 rpm. The digestion was stopped with 2% formic acid (FA) and precipitated DOC was removed by filtration using 0.2 μm PVDF filters.

### AP-MS

Cell pellets were lysed by resolubilizing in 2 ml HNN buffer supplemented with 0.5% IGEPAL or 1% DDM when using PRKCA as bait, 400 nM Na_3_VO_4_, 1 mM PMSF, 0.2% Protease Inhibitor Cocktail (Sigma) and 0.5 μl/ml benzonase (Sigma). The lysate was incubated on ice for 10 minutes and centrifuged at 18000x g for 20 minutes at 4°C. The pulldown was done as for AP-LiP-MS samples, except that the final wash was done with 100 mM ammonium bicarbonate. Samples were eluted twice by incubation with 30 μl of 0.5 mM biotin in 100 mM ammonium bicarbonate for 15 minutes. To the samples, 60 μl 8 M urea in 100 mM ammonium bicarbonate for a final concentration of 4 M urea. Samples were reduced and alkylated as for AP-LiP-MS. Samples were diluted with 100 mM ammonium bicarbonate to a urea concentration of 1 M. To each sample, 1 μg Lys-C and 1 μg trypsin was added and samples were incubated o/n at 37°C and 200 rpm. The digestion was stopped with 2% FA.

### *In vivo* proximity labeling

Cell pellets were lysed in 1 ml RIPA buffer (50 mM Tris-HCl pH 8, 150 mM NaCl, 1 % Triton, 1mM EDTA, 0.1 % SDS) supplemented with 1 mM PMSF, 0.2% Protease Inhibitor Cocktail (Sigma) and 0.5 μl/ml benzonase (Sigma). The lysate was briefly sonicated for 30 s, incubated at 10°C for 30 minutes and centrifuged at 18000x g for 20 minutes at 4°C. The supernatant was added to protease resistant 120 μl StrepTactin Sepharose slurry (IBA Lifesciences) that were modified as previously described^97^ and incubated on a rotating wheel for 60 minutes at 4°C. The beads were loaded onto a 1 μm glass filter plate and washed 3 times with 1 ml RIPA buffer, 3 times with 1 ml HNN buffer, 3 times with 100 mM ammonium bicarbonate and transferred to a 10 kDa cutoff plate and centrifuged at 1500x g for 15 minutes. To the samples, 100 μl of 8 M urea was added. Samples were reduced with 10 mM TCEP for 30 minutes at 37 °C and 200 rpm, and alkylated with 40 mM IAA for 30 minutes at 37°C in the dark, at 200 rpm. Following centrifugation at 1500x g for 15 minutes, beads were washed twice with 250 μl of 100 mM ammonium bicarbonate and resuspended in 200 μl of 100 mM ammonium bicarbonate. Samples were digested o/n at 37°C at 200 rpm in the presence of 1 μg trypsin and 1 μg Lys-C. Peptides were collected by centrifugation at 1500x g for 20 minutes from the supernatant and combined with the peptides from the supernatant obtained from washing the digested beads with 100 μl of 100 mM ammonium bicarbonate and centrifugation at 1500x g for 40 minutes. The digestion was stopped with 5% FA.

### Desalting

Samples were desalted using HNFR S18V desalting plates (Nest Group). The resin was activated using 200 μl methanol and washed 3 times with 200 μl Buffer B (50% ACN and 0.1% FA for AP-LiP and AP samples, 40% ACN and 0.1% FA for proximity labeling samples). The resin was equilibrated 3 times with 200 μl Buffer A (0.1% FA for AP-LiP and AP samples, 5% ACN and 0.1% FA for proximity labeling samples). Samples were loaded and washed 3 times with 200 μl Buffer A. Samples were eluted twice with 100 μl Buffer B and dried in a vacuum centrifuge.

### Mass spectrometry

AP-LiP-MS, AP-MS and PRKCA proximity labeling experiments were measured on a Orbitrap Fusion Lumos Tribrid mass spectrometer (Thermo Fisher Scientific) equipped with an Acquity M-Class UPLC System (Waters). To separate the peptides on self-pack 40 cm x 0.75 μm fritted emitter (CoAnn) packed with 3 μm C18 beads (Dr. Maisch), we used a linear gradient of 3% to 35% acetonitrile in water with 0.1% FA over the course of 120 minutes at flow rate of 300 nl/min. Library samples consisting of pools of replicates were measured by DDA. The MS1 spectra were acquired in a scan range of 350-1150m/z with an Orbitrap resolution of 120000. The normalized automatic gain control (AGC) target was set to 200% with a maximum injection time of 100 ms. The RF lens was set to 30%. The targeted MS2 spectra were acquired for the top 20 peaks in the spectrum and fragmented with a HCD of 30%. The spectra were measured with an Orbitrap resolution of 30000 with isolation windows of 1.6 m/z. The AGC target was set to 200% with a maximum injection time of 54 ms. Samples for quantification were measured by DIA. The MS1 spectra were acquired in a scan range of 350-1400 m/z with an Orbitrap resolution of 120000. The normalized AGC target was set to 50% with a maximum injection time of 100 ms. The RF lens was set to 30%. The targeted MS2 spectra were acquired for the desired masses with variable isolation windows (Supplementary Table 5) and fragmented with a HCD of 28%. The spectra were measured with an Orbitrap resolution of 30000 with variable scan ranges. The AGC target was set to 200% with a maximum injection time of 54 ms. The RF lens was set to 30%.

Proximity labeling samples were measured on an Orbitrap Exploris 480 mass spectrometer (Thermo Fischer Scientific) equipped with an EASY-nLC 1000 System (Thermo Fischer Scientific). Peptides were separated on the same column used for the Orbitrap Fusion Lumos Tribrid mass spectrometer using a linear gradient of 5% to 30% acetonitrile in water (0.1% FA) over the course of 120 minutes at flow rate of 300 nl/min. Library samples consisting of pools of replicates were measured by DDA. The MS spectra were acquired at an Orbitrap resolution of 120000 between 350 m/z and 1500 m/z. The normalized AGC target was set to 200% or a maximum injection time of 100 ms. Subsequently, 20 MS/MS spectra were acquired after higher-energy collisional dissociation (HCD) with a normalized collision energy of 30%, an Orbitrap resolution of 30000, and a normalized AGC target of 200% or a maximum injection time of 54 ms. Precursors with a minimum intensity of 5000 ion counts and a charge state between 2 and 6 were selected with a mass window of 1.4 m/z. A dynamic exclusion window of 20 s was used.

Samples for quantification were measured by DIA. The MS1 spectra were acquired in a scan range of 350-1150m/z with an Orbitrap resolution of 120000. The normalized AGC target was set to 200% with a maximum injection time of 264 ms. The RF lens was set to 50%. The targeted MS2 spectra were acquired for the desired masses with variable isolation windows (Supplementary Table 6) and fragmented with a HCD of 30%. The spectra are measured with an Orbitrap resolution of 30000 with variable scan ranges. The AGC target was set to 200% with a maximum injection time of 66 ms. The RF lens was set to 50%.

### Search of Mass Spectrometry data

Spectra were searched using Spectronaut v. 19 in directDIA mode with the measured DDA library files used to extend the library. Searches were done separately for each experiment and bait. For AP-LiP-MS and AP-MS data, semi-specific peptides were included in the search, and phosphorylation was included as a variable modification. For proximity labeling data, only specific peptides were included in the search, and the number of missed cleavages allowed was four. Both biotinylation and phosphorylation were included as variable modifications. For all searches, cross-run normalization was turned off. Report files were exported and imported into R version 4 and analyzed using protti version 0.6.0^98^.

### Determination of structural changes in AP-LiP-MS data

The code for all analyses is available on github.com/vivreb/multi-proteomic_profiling_of_kinases/. Identified peaks were filtered for q-value < 0.001, log_2_(intensity) > 10 and Δrt < 0.1 minutes, where Δrt was defined as the difference between the predicted retention time and the empirical apex retention time. Peptides were filtered for proteotypicity and common contaminants were removed. Only peptides that were complete across all conditions were used for further analysis. Peptide abundance was normalized based on rank invariant values using the MBQN package^99^. Differential abundance was calculated using a moderated t-test^100^. Multiple testing correction was performed using the Benjamini-Hochberg procedure^101^. Peptides with an absolute log_2_(fold change) > 0.585 (corresponding to a fold change > 1.5) and adjusted p-value < 0.05 were considered significantly changing.

### Determination of interactors in AP-MS data

Identified peaks were filtered for q-value < 0.001, log_2_(intensity) > 9 and Δrt < 1 minute. Peptides were filtered for specificity and proteotypicity. Only peptides that were found in all replicates within a given condition were used for further analysis. Peptide abundance was normalized based on rank invariant values using the MBQN package^99^ and bait normalized. Protein abundance was calculated by MaxLFQ^102^ for proteins with at least 3 peptides. We used a modified version of the weighted D-score^103^ to determine interactors. First, we used protein intensity instead of spectral count, and we augmented the D-score 1.1-fold for known interactors based on physical interactions in the BioGrid database^38^. We consider proteins that have a modified D-score > 1.425 as interactors (Supplementary Fig. 10A). We confirmed that we effectively recover known interactors using a ROC area under the curve (AUC) analysis (AUC = 0.992, Supplementary Fig. 10B).

### Determination of interaction changes in AP-MS data

Peptides were filtered and normalized as for the determination of interactors before imputation using msImpute^104^ and protein abundance calculation using MaxLFQ^102^. For all interactors, the differential abundance was calculated using a moderated t-test^100^. Multiple testing correction was performed using the Benjamini-Hochberg procedure^101^. Proteins with an absolute log_2_(fold change) > 1 and adjusted p-value < 0.05 were considered significantly changing.

### Determination of phosphorylation changes in AP-MS data

Filtering was done as for the AP-LiP-MS analysis. Peptide abundance was normalized based on rank invariant values using the MBQN package^99^ and bait normalized. Only peptides complete in at least one condition were kept for further analysis, and peptides were imputed using msImpute^104^. Only peptides containing a phosphorylation were considered for the analysis. Differential abundance was calculated using a moderated t-test^100^. Multiple testing correction was performed using the Benjamini- Hochberg procedure^101^. Peptides with an absolute log_2_(fold change) > 0.585 (corresponding to a fold change > 1.5) and adjusted p-value < 0.05 were considered significantly changing.

### Determination of proximity changes in proximity labeling data

Identified peaks were filtered for q-value < 0.0001, log_2_(intensity) > 9 and Δrt < 0.5 minutes. Peptides were filtered for proteotypicity and common contaminants were removed. Peptide abundance was normalized based on rank invariant values using the MBQN package^99^. Protein abundance was calculated by MaxLFQ^102^ for proteins with at least 3 peptides. Differential abundance was calculated using a moderated t-test^100^. Multiple testing correction was performed using the Benjamini-Hochberg procedure^101^. The same calculations were performed for an EGFP control, and proteins showing a change in the same direction as in the control were removed from the data. Adjusted p-values were recalculated for the revised dataset. Proteins with an absolute log_2_(fold change) > 1 and adjusted p-value < 0.05 were considered significantly changing.

### siRNA-mediated knockdown

HEK293 cells (ATCC) were seeded at 10^5^ cells per well in a 6-well plate in DMEM supplemented with 10% dialyzed FBS (BioConcept). To each well, 50 μM siRNA (siCAMKK2: s20926, Thermo Fisher Scientific; siNEG: iSilencer/i Negative Control No. 1, Thermo Fisher Scientific) and 7 μl siPORT NeoFX transfection reagent (Thermo Fisher Scientific) was added. Cells were grown for 72 h. Samples were treated with 10 μM SGC-CAMKK2-1 for 4 h in 0.01% DMSO and glucose starved for 20 min in glucose-free DMEM (Gibco) supplemented with 10% dialyzed FBS (BioConcept) before harvesting. Samples were harvested in 100 μl RIPA buffer supplemented with PI, PMSF, Na3VO4 and 0.5μg/ml benzonase and snap frozen. Samples were stored at -80°C until processing.

### Western blot analysis

Samples were thawed on ice, and centrifuged at 18000x g for 20 minutes and the supernatant was collected. Total protein concentration was measured using a BCA assay (Pierce), and the samples were diluted to 2.5 μg/μl. The samples were mixed with 4x Lämmli buffer (Invitrogen) and heated to 95°C for 5 minutes. Of the samples, 20 µl were loaded onto pre-cast 4-12% Bis-Tris gels (Invitrogen) along with 5 µl of the ladder (BioRad). The gel was run at 100 V for 60 minutes. The gel was blotted onto a nitrocellulose membrane (BioRad) for 30 minutes. The membrane was blocked with 5% milk in TBST (20 mM Tris pH 7.6, 150 mM NaCl, 1% TWEEN 20) for 1 h. The primary antibody (anti-pT183 PRKAA1: Cell Signaling Technology (D4D6D); anti-PRKAA1: Cell Signaling Technology (2532); anti-CAMKK2: Proteintech (11549-1-AP); anti-γH2AX: Merck (05-636); anti-H2AX: Sigma (PLA0023); anti-GAPDH: Invitrogen (GA1R)) was added for 2 h at room temperature or o/n at 4°C and subsequently washed for 3x 10 minutes with TBST. The secondary antibody (anti-mouse secondary antibody: Cell Signaling Technology (7076S), anti-rabbit secondary antibody: Cell Signaling Technology (7074S); all diluted 1:1000) was added for 1 hour at RT. The membrane was again washed for 3x 10 minutes. The ECL detection reagent (Cytiva) was added and the membrane was imaged by chemiluminescence with various exposure times. The antibodies were removed by incubation in western blot stripping buffer (Invitrogen) for 10 min, washed in TBST and blocked before the next antibody was added.

### AlphaFold3 Prediction and Structural Analysis

The AlphaFold3^47^ prediction of the CAMKK2-PRKAA1 complex was generated using the AlphaFold3 webserver (alphafoldserver.com/). Only the sequences of the KDs were used (Supplementary Table 7). The interface analysis was done in the PyMOL Molecular Graphics System version 2.5.3. The interface comprises residues that are within 2.8 Å of the other protein and show a change in accessible surface area (ΔASA) of at least 1 Å^2^ upon binding the interaction partner.

### Live cell imaging

HEK293 cells expressing N-terminally tagged EGFP-PRKCA were seeded at 10^5^ cells in 35 mm glass-bottom cell culture dishes (Ibidi) in DMEM supplemented with 10% dialyzed FBS (BioConcept) and 1% P/S, and induced for 24 h with 1.3 μg/ml doxycycline. For imaging, FluoroBrite DMEM (Gibco) supplemented with 10% dialyzed FBS (BioConcept), 1% P/S and 1.3 μg/ml doxycycline was used. To the cells, 1 μM Gö 6983 in 0.01% DMSO diluted in 100 μl imaging medium was added and cells were immediately imaged using an instant Structured Illumination Microscope (VisiTech). Z-stacks were acquired across 5 time points 2 minutes apart. Images were deconvolved using Huygens Professional 24.10. Individual slices from representative cells were normalized and cropped using ImageJ and a scale bar showing 5 μm was added.

### Immunofluorescence microscopy

HeLa CCL2 cells (ATCC) were seeded at 10^5^ cells per well in a 6-well plate containing cover slips in DMEM supplemented with 10% dialyzed FBS (BioConcept) and 1% P/S and grown o/n. To the cells, 1 μM rabusertib for 4 h, 6 μM RO-3306 (Sigma) for 24 h, or 50 μM etoposide (Sigma) for 3 h was added in 0.01% DMSO. The cells were incubated with 100 nM MitoTracker Deep Red FM (Invitrogen) for 1 h. Cells were subsequently washed with warm PBS and fixed with 4% PFA in PBS (Invitrogen) for 15 minutes. Cells were washed and the coverslips mounted on slides using ProLong Diamond mounting medium (Invitrogen). Z-stacks of the samples were imaged using an instant Structured Illumination Microscope (VisiTech). Images were deconvolved using Huygens Professional 24.10 and cropped using ImageJ. Image segmentation was performed in Ilastik 1^105^. The number of pixels from fragmented mitochondria were counted and normalized based on the total number of pixels from mitochondria in each image. Significance was calculated per experiment. To assess significance of rabusertib treatment with and without RO-3306 treatment, we used two-way analysis of variance (ANOVA) followed by a Tukey Honestly Significant Difference (HSD) test. To separately test the effects of etoposide treatment, we used a two-tailed t-test. 10 images taken from 3 replicates were quantified for each condition.

### Fluo-4 calcium imaging

HEK293 cells ectopically expressing miniTurbo-PRKCA were grown in DMEM supplemented with 10% dialyzed FBS (BioConcept) and 1% P/S, induced with 1.3 μg/ml doxycycline and treated with 1 μM Gö 6983 or 0.01% DMSO for 4 h. Fluo-4 solution was prepared according to manufacturer’s instructions (Invitrogen). Baseline fluorescence was measured with an excitation wavelength of 483 nM and an emission wavelength of 530 nM every 58 seconds for 10 cycles. To the samples, 10 μM ionomycin or 1% ethanol, or 1 μM Gö 6983 or 1% DMSO was added. After addition of the reagents, the measurements were repeated for 250 additional cycles. Measurements were normalized to a baseline control without the Fluo-4 reagent and to the respective vehicle control. All measurements were done in six replicates and mean and standard error of the mean were reported.

## Data and code availability

All MS data and related files are accessible via the ProteomeXchange Consortium^106^ via the PRIDE partner repository with the dataset identifier PXD069830. All code and additional data necessary to fully reproduce all analyses is available for download on github via github.com/vivreb/multi-proteomic_profiling_of_kinases/.

## Disclosure and competing interest statement

P.P. is an inventor on a patent that covers the LiP-MS method, a component of the pipeline presented in this manuscript. The remaining authors declare no competing interests.

## Acknowledgements

We thank the group of Dr. Anne-Claude Gingras for the pcDNA5-pDEST-FRT-MiniTurbo-Flag vectors. We thank Dr. Natalie de Souza for her comments on the manuscript. We thank ScopeM for their support and assistance with microscopy experiments. This work was supported by the Innovative Health Initiative 2 Joint Undertaking (JU) under grant agreement No 875510. The JU receives support from the European Union’s Horizon 2020 research and innovation program and EFPIA and Ontario Institute for Cancer Research, Royal Institution for the Advancement of Learning McGill University, Kungliga Tekniska Hoegskolan, Diamond Light Source Limited.

## Notes

https://www.github.com/vivreb/multi-proteomic_profiling_of_kinases/

### Summary of Updates

Results and analyses remain the same, only framing was altered

