## Supplementary information for "Paradoxical non-catalytic kinase functions are driven by inhibitor-induced displacement of autoinhibitory domains"

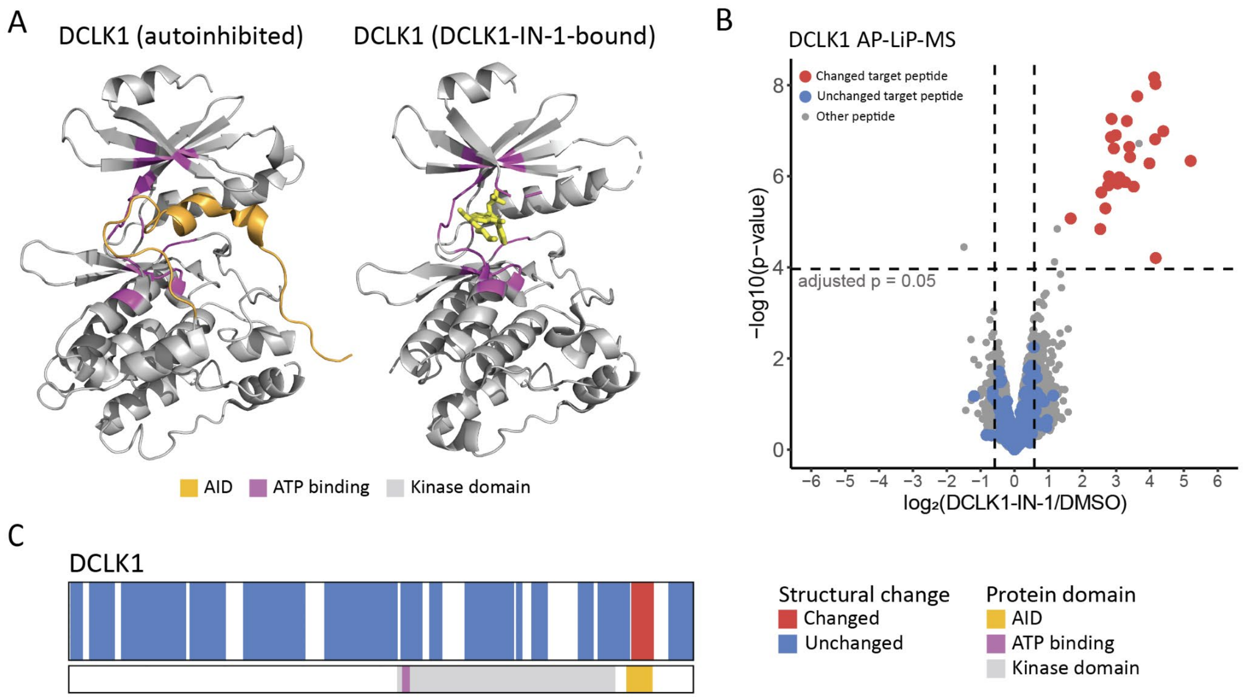


**Supplementary Fig. 1. Validation of AP-LiP-MS using DCLK1 inhibited with DCLK1-IN-1.** A) Comparison of the resolved structures of DCLK1 in the autoinhibited configuration (PDB: 6KYQ) and the DCLK1-IN-1 bound configuration (PDB: 7KXW; the inhibitor DCLK1-IN-1 is shown in yellow). B) Volcano plot showing LiP peptide intensity changes in the DCLK1-enriched samples upon treatment with DCLK1-IN-1 compared to vehicle control. In red are significantly changing DCLK1 peptides, in blue are the non-changing DCLK1 peptides. Peptides from other proteins are shown in grey. C) Barcode plot of structural changes in DCLK1 in response to inhibition with DCLK1-IN-1.

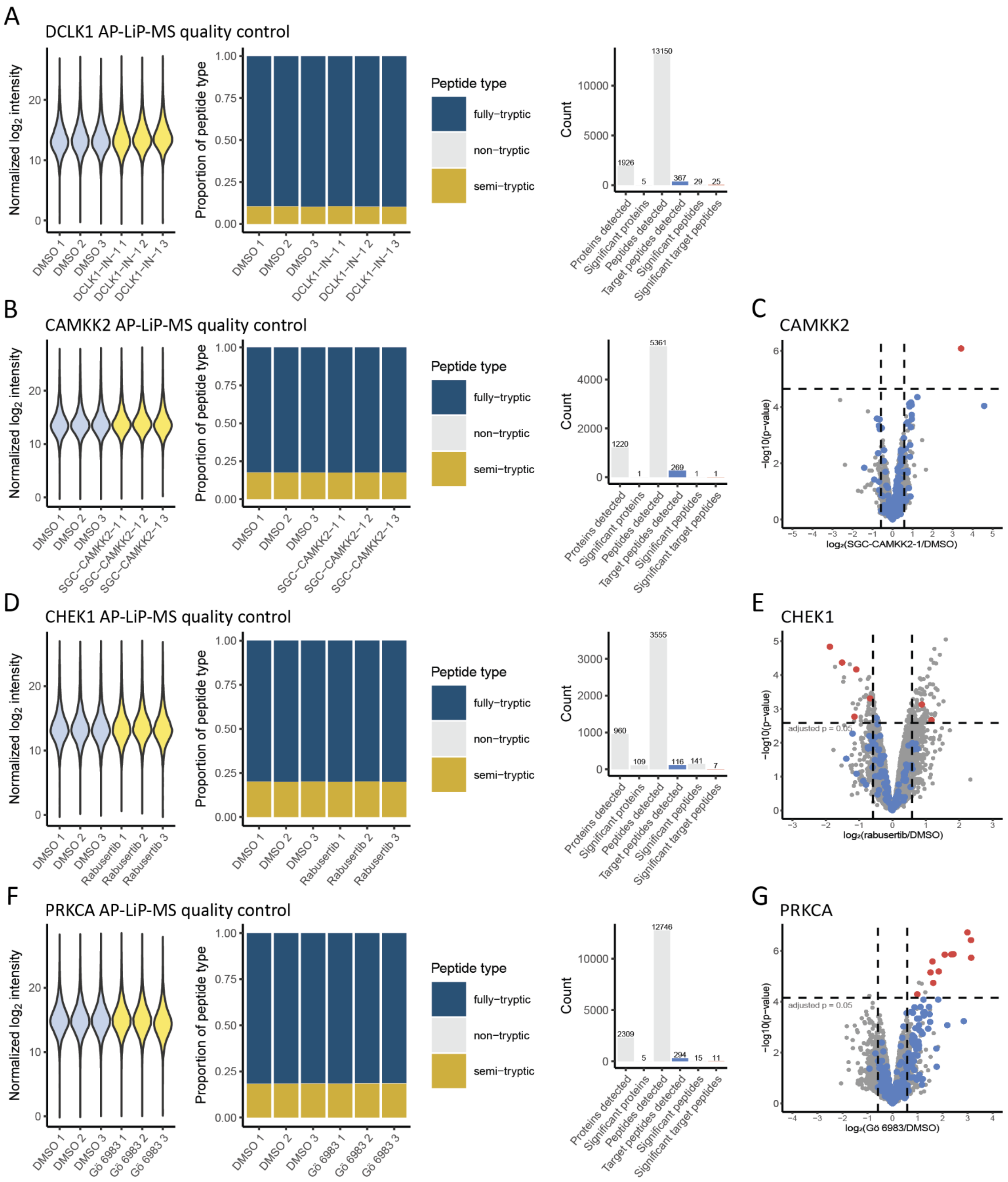


**Supplementary Fig. 2. Quality control for AP-LiP-MS data.** A, B, D, F) Normalized raw intensities, raw peptide type distribution, and summary of metrics related to the peptides quantified for DCLK1, CAMKK2, CHEK1 and PRKCA, respectively. C, E, G) Volcano plot showing LiP peptide intensity changes in the kinase-enriched samples for CAMKK2, CHEK1, and PRKCA, respectively, upon treatment with SGC-CAMKK2-1, rabusertib, or Gö 6983, compared to vehicle control. In red are significantly changing target peptides, in blue are the non-changing target peptides. Peptides from other proteins are shown in grey. All experiments were performed in triplicates and p-values were calculated using a moderated t-test followed by multiple testing corrections using the procedure of Benjamini-Hochberg.

**Supplementary Table 1: Interactors of CAMKK2 identified by AP-MS across all treatment conditions.** Interactors were determined by bait and treatment condition using a modified version of the weighted D-score (see Methods for details). Any protein with a score above 1.425 was considered an interactor.

| Gene name | UniProt Accession | Known BioGrid Interactor | Maximal score |
|---|---|---|---|
| ACAA2 | P42765 | FALSE | 1.44024 |
| AFG3L2 | Q9Y4W6 | FALSE | 1.473367 |
| ALDH3A2 | P51648 | FALSE | 1.46762 |
| ARAF | P10398 | FALSE | 1.456088 |
| ATP1B3 | P54709 | FALSE | 1.437549 |
| CAMKK2 | Q96RR4 | TRUE | 1.745324 |
| CDC37 | Q16543 | TRUE | 1.518405 |
| CDC42EP1 | Q00587 | FALSE | 1.490695 |
| COPB1 | P53618 | FALSE | 1.463998 |
| CTSB | P07858 | FALSE | 1.445539 |
| DCAF8 | Q5TAQ9 | FALSE | 1.46323 |
| DOP1B | Q9Y3R5 | FALSE | 1.448148 |
| DYNC1LI2 | O43237 | FALSE | 1.450658 |
| ECPAS | Q5VYK3 | FALSE | 1.442603 |
| ESYT2 | A0FGR8 | TRUE | 1.562158 |
| FAM83H | Q6ZRV2 | FALSE | 1.476072 |
| FANCD2 | Q9BXW9 | FALSE | 1.427346 |
| FANCI | Q9NVI1 | TRUE | 1.481436 |
| FGF2 | P09038 | FALSE | 1.430508 |
| FLNA | P21333 | TRUE | 1.45766 |
| FOXK2 | Q01167 | FALSE | 1.432111 |
| GCN1 | Q92616 | TRUE | 1.500203 |
| GGCX | P38435 | FALSE | 1.456247 |
| HIP1R | O75146 | FALSE | 1.441217 |
| HK1 | P19367 | FALSE | 1.494657 |
| HSP90AA1 | P07900 | TRUE | 1.494427 |
| HSPA2 | P54652 | FALSE | 1.465704 |
| HSPA4L | O95757 | TRUE | 1.471913 |
| IQGAP2 | Q13576 | TRUE | 1.467299 |
| IRAK1 | P51617 | TRUE | 1.482286 |
| KNTC1 | P50748 | FALSE | 1.445306 |
| MARK4 | Q96L34 | FALSE | 1.456161 |
| MON2 | Q7Z3U7 | FALSE | 1.474501 |
| MYO6 | Q9UM54 | FALSE | 1.445281 |
| NDUFS7 | O75251 | FALSE | 1.479526 |
| NF1 | P21359 | FALSE | 1.426087 |
| NPEPPS | P55786 | FALSE | 1.43914 |
| OBSL1 | O75147 | TRUE | 1.531543 |
| PARK7 | Q99497 | TRUE | 1.457196 |
| PFKM | P08237 | FALSE | 1.473358 |
| PPP6C | O00743 | FALSE | 1.467793 |
| PPP6R3 | Q5H9R7 | FALSE | 1.471 |
| PRKAA1 | Q13131 | TRUE | 1.655716 |
| PRKAA2 | P54646 | TRUE | 1.62695 |

| | | | |
|---|---|---|---|
| PRKAG1 | P54619 | TRUE | 1.671049 |
| PRKDC | P78527 | TRUE | 1.471861 |
| SCO2 | O43819 | FALSE | 1.466272 |
| SLC25A1 | P53007 | TRUE | 1.471115 |
| SLC25A11 | Q02978 | TRUE | 1.490104 |
| SLC3A2 | P08195 | FALSE | 1.453671 |
| SMC1A | Q14683 | TRUE | 1.464657 |
| SMC3 | Q9UQE7 | TRUE | 1.459327 |
| SPTLC1 | O15269 | FALSE | 1.465125 |
| STON2 | Q8WXE9 | FALSE | 1.464973 |
| TANGO6 | Q9C0B7 | FALSE | 1.445534 |
| THAP11 | Q96EK4 | TRUE | 1.483681 |
| TM9SF3 | Q9HD45 | FALSE | 1.466035 |
| TMEM209 | Q96SK2 | FALSE | 1.444921 |
| TMEM263 | Q8WUH6 | FALSE | 1.468814 |
| TUBGCP2 | Q9BSJ2 | FALSE | 1.482299 |
| UBE3C | Q15386 | FALSE | 1.449353 |
| VPS4A | Q9UN37 | FALSE | 1.445089 |
| XPOT | O43592 | TRUE | 1.488375 |
| ZMAT5 | Q9UDW3 | FALSE | 1.475343 |
| ZNF174 | Q15697 | FALSE | 1.478126 |
| ZNF92 | Q03936 | FALSE | 1.493794 |

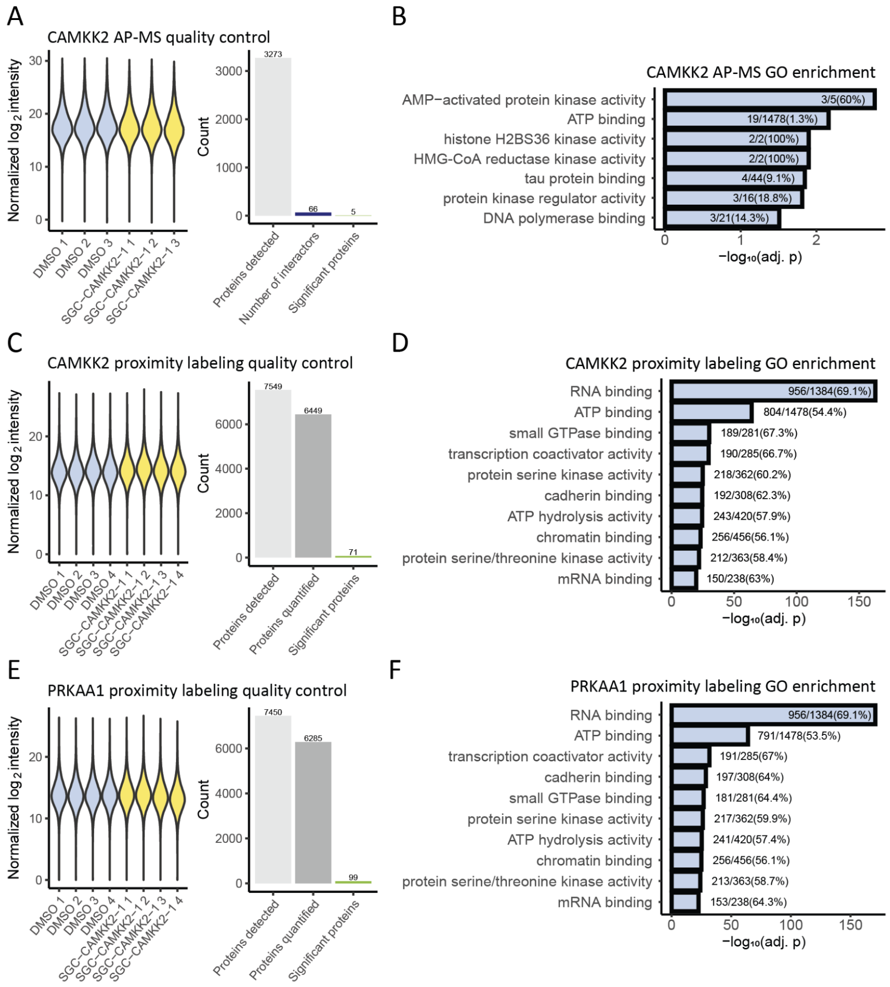


**Supplementary Fig. 3. Quality control for characterization of CAMKK2 complexes and proximity upon SGC-CAMKK2-1 treatment.** A, C, E) Normalized raw intensities and summary of metrics related to the proteins quantified for CAMKK2 AP-MS, CAMKK2 *in vivo* proximity labeling, and PRKAA1 *in vivo* proximity labeling, respectively. B, D, F) GO enrichment analysis (molecular function) across all CAMKK2 interactors identified in the AP-MS experiment, or all proteins quantified in the *in vivo* proximity labeling experiment using CAMKK2 or PRKAA1 as bait, respectively. P-values were calculated using a Fisher's exact test followed by multiple testing corrections using the procedure of Benjamini-Hochberg.

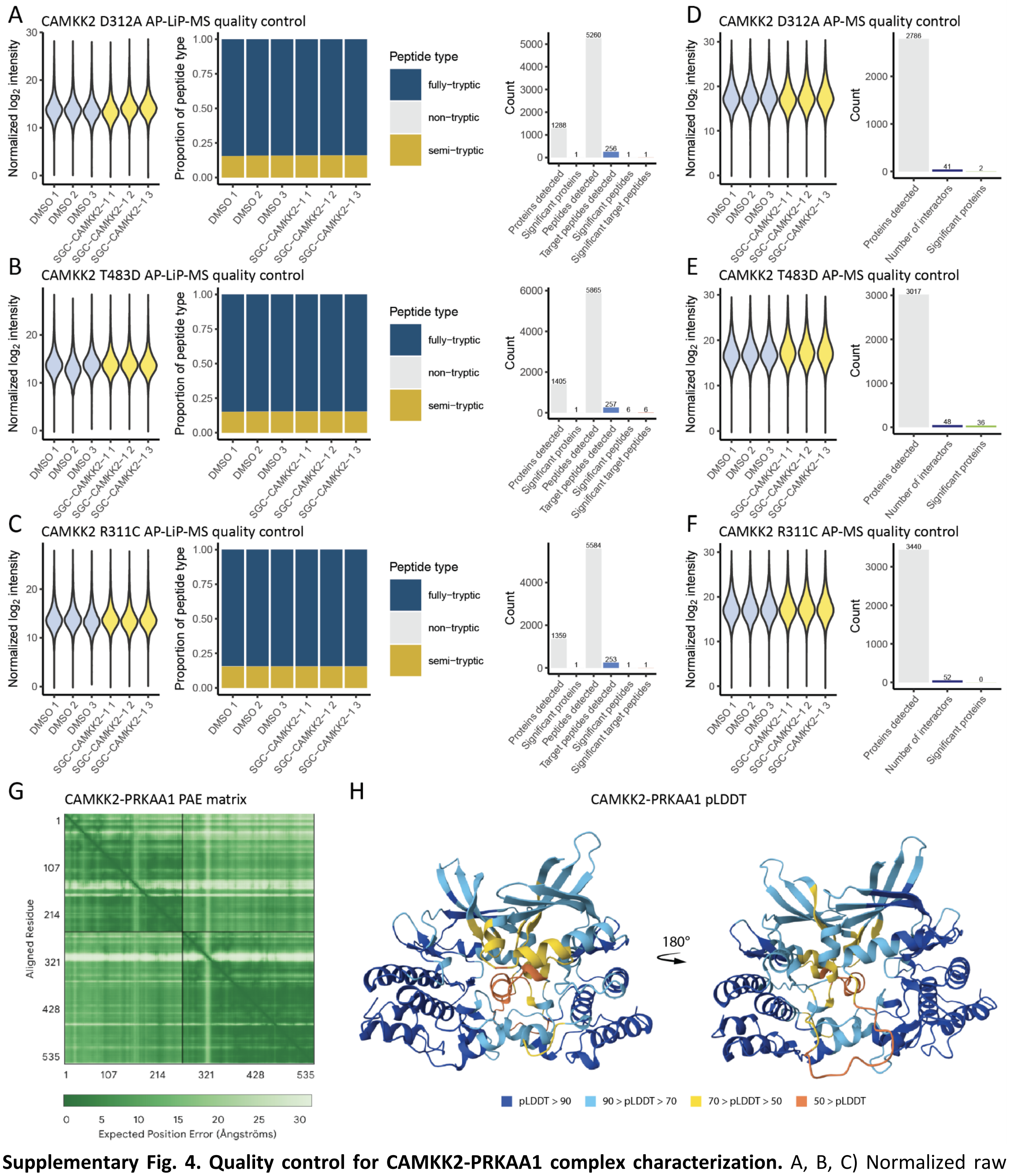


**Supplementary Fig. 4. Quality control for CAMKK2-PRKAA1 complex characterization.** A, B, C) Normalized raw intensities, raw peptide type distribution, and summary of metrics related to the peptides quantified for AP-LiP-MS to analyze changes in CAMKK2 D312A upon SGC-CAMKK2-1 treatment. D, E, F) Normalized raw intensities and summary of metrics related to the proteins quantified for CAMKK2 D312A, T483D and R331C AP-MS, respectively. G) Predicted Aligned Error (PAE) matrix for the CAMKK2-PRKAA1 AlphaFold3 prediction. H) The predicted local distance difference test (pLDDT) score mapped onto the predicted structure of the CAMKK2-PRKAA1 structure from two different views. AP-LiP-MS and AP-MS experiments were performed in triplicates.

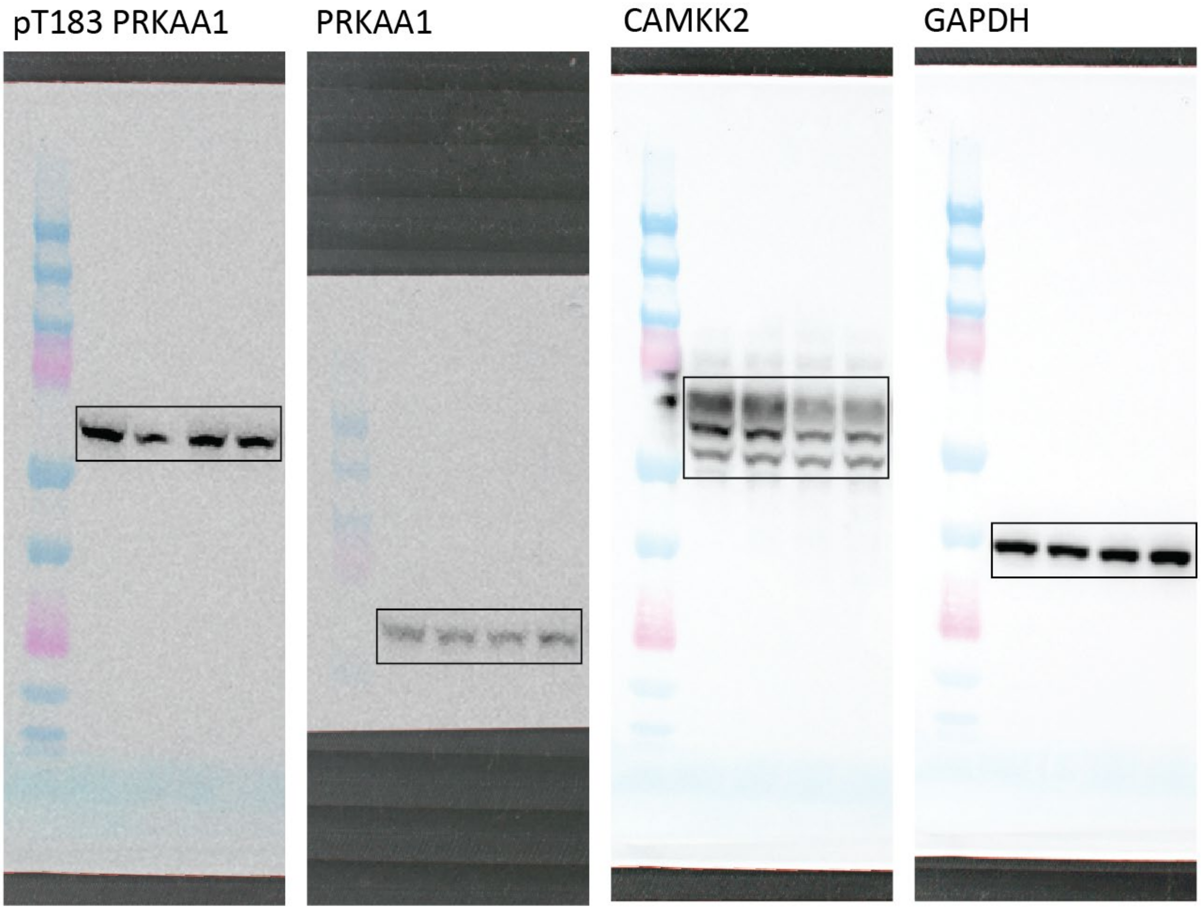


**Supplementary Fig. 5. Raw western blot images acquired for Fig. 3F.** The boxes show the cropped areas used in the final images.

**Supplementary Table 2: Interactors of CHEK1 identified by AP-MS across all treatment conditions.** Interactors were determined by bait and treatment condition using a modified version of the weighted D-score (see Methods for details). Any protein with a score above 1.425 was considered an interactor.

| Gene name | UniProt Accession | Known BioGrid Interactor | Maximal score |
|---|---|---|---|
| BLM | P54132 | TRUE | 1.449943 |
| CAPZB | P47756 | TRUE | 1.454838 |
| CCDC12 | Q8WUD4 | FALSE | 1.480635 |
| CDC37 | Q16543 | TRUE | 1.486203 |
| CDC5L | Q99459 | TRUE | 1.476245 |
| CDK1 | P06493 | TRUE | 1.472145 |
| CHEK1 | O14757 | TRUE | 1.664952 |
| CLPB | Q9H078 | TRUE | 1.615227 |
| CSNK2B | P67870 | TRUE | 1.456171 |
| CTBP1 | Q13363 | FALSE | 1.464498 |
| CUL4A | Q13619 | TRUE | 1.479329 |
| CWC22 | Q9HCG8 | FALSE | 1.431847 |
| DCAF8 | Q5TAQ9 | FALSE | 1.425562 |
| DDB1 | Q16531 | TRUE | 1.491276 |
| FUS | P35637 | TRUE | 1.454455 |
| H1-10 | Q92522 | TRUE | 1.513278 |
| HMGB3 | O15347 | FALSE | 1.446154 |
| HSP90AA1 | P07900 | TRUE | 1.488287 |
| LSM3 | P62310 | FALSE | 1.483499 |
| MCM2 | P49736 | TRUE | 1.449287 |
| MCM3 | P25205 | TRUE | 1.485414 |
| MCM4 | P33991 | TRUE | 1.438203 |
| MCM5 | P33992 | TRUE | 1.448998 |
| MCM6 | Q14566 | TRUE | 1.455789 |
| MEN1 | O00255 | TRUE | 1.472779 |
| MSH6 | P52701 | TRUE | 1.442098 |
| NPM1 | P06748 | TRUE | 1.499068 |
| NPM3 | O75607 | FALSE | 1.440408 |
| PARP1 | P09874 | TRUE | 1.485756 |
| PCNA | P12004 | TRUE | 1.464558 |
| PM20D2 | Q8IYS1 | FALSE | 1.440793 |
| PPM1G | O15355 | FALSE | 1.428902 |
| PRKDC | P78527 | TRUE | 1.461032 |
| PSMD4 | P55036 | FALSE | 1.451336 |
| RACGAP1 | Q9H0H5 | TRUE | 1.456254 |
| RPS27A | P62979 | TRUE | 1.521202 |
| SNW1 | Q13573 | TRUE | 1.458479 |
| SRM | P19623 | FALSE | 1.472898 |
| TOM1 | O60784 | TRUE | 1.467789 |
| TP53 | P04637 | TRUE | 1.457303 |
| UBA1 | P22314 | TRUE | 1.434492 |

| | | | |
|---|---|---|---|
| USP7 | Q93009 | TRUE | 1.460129 |
| VCP | P55072 | TRUE | 1.453499 |
| WDR48 | Q8TAF3 | FALSE | 1.439921 |
| XRCC5 | P13010 | TRUE | 1.469176 |
| XRCC6 | P12956 | TRUE | 1.463784 |
| YWHAB | P31946 | TRUE | 1.47423 |
| YWHAG | P61981 | TRUE | 1.460321 |
| YWHAH | Q04917 | TRUE | 1.459052 |
| YWHAQ | P27348 | TRUE | 1.467858 |
| YWHAZ | P63104 | TRUE | 1.460007 |

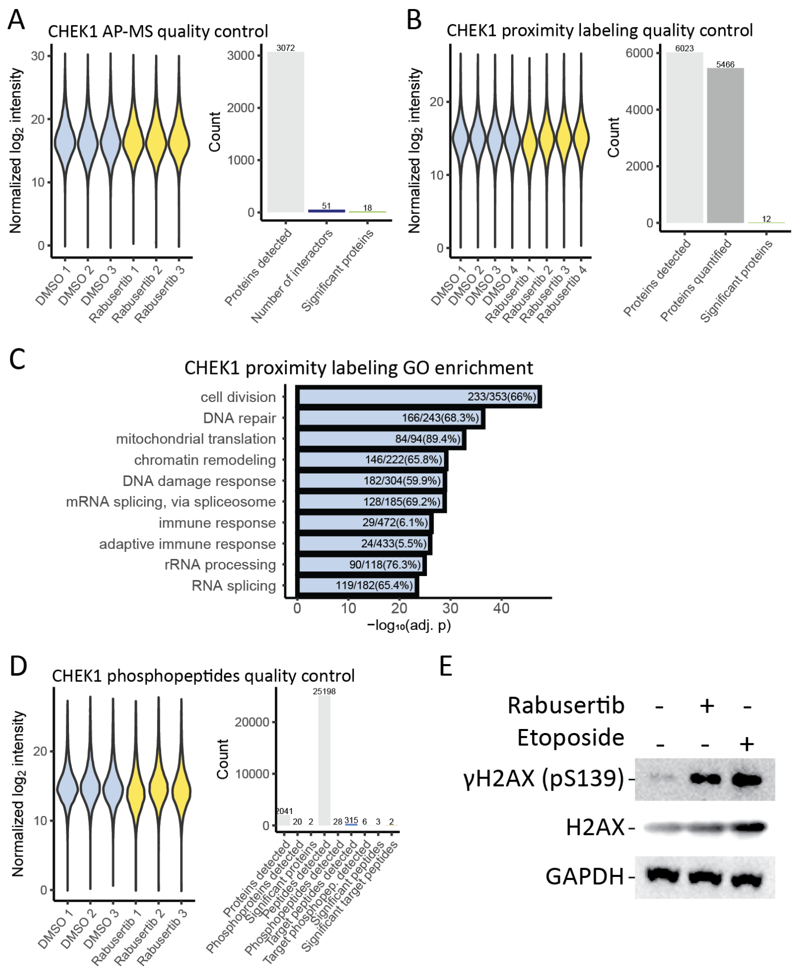


**Supplementary Fig. 6. Quality control for characterization of CHEK1 complexes and proximity upon rabusertib treatment.** A, B) Normalized raw intensities and summary of metrics related to the proteins quantified for CHEK1 AP-MS and *in vivo* proximity labeling, respectively. C) GO enrichment analysis (biological process) across all proteins quantified in the *in vivo* proximity labeling experiment using CHEK1 as bait. Only enriched terms were plotted. P-values were calculated using a Fisher's exact test followed by multiple testing corrections using the procedure of Benjamini-Hochberg. D) Normalized raw intensities and summary of metrics related to the peptides quantified for CHEK1 phosphopeptide analysis. E) Western blot analysis of DNA damage in HEK cells upon treatment with rabusertib (4h) or etoposide (3h). AP-MS experiments were performed in triplicates, and *in vivo* proximity labeling experiments in quadruplicates.

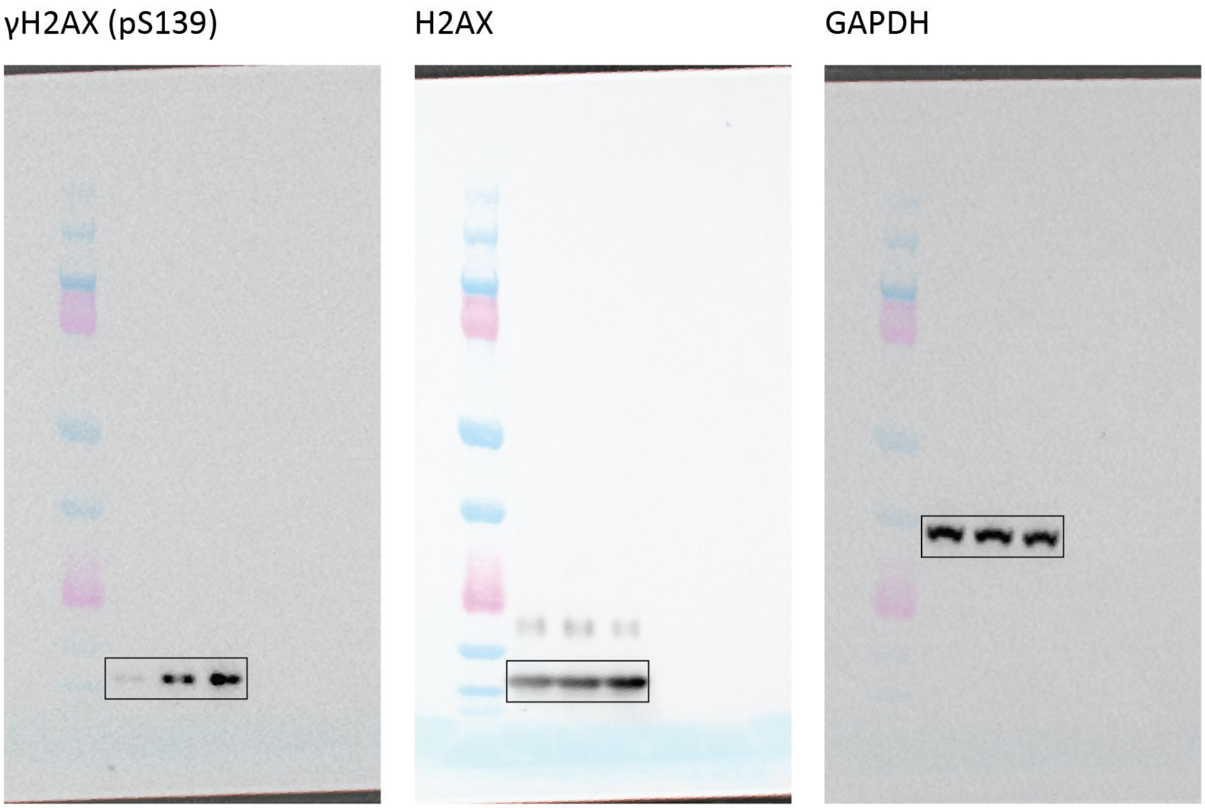


**Supplementary Fig. 7. Raw western blot images acquired for Supplementary Fig. 6E.** The boxes show the cropped areas used in the final images.

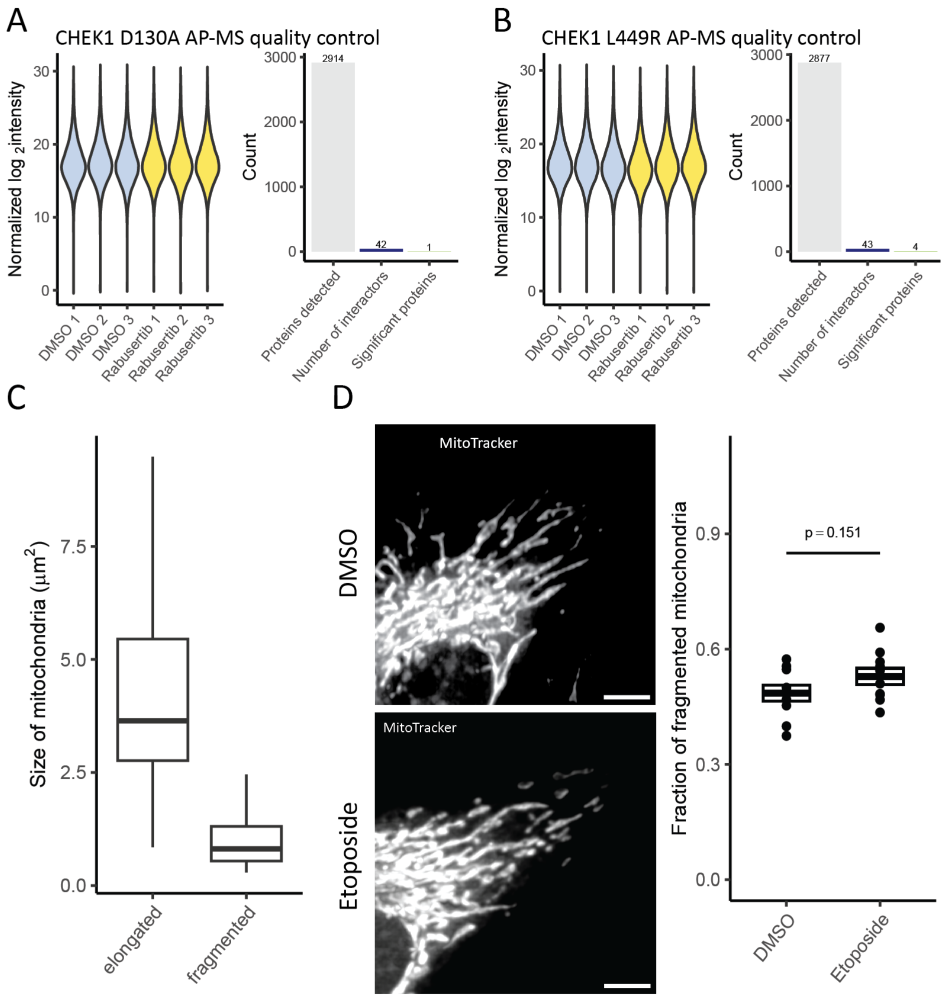


**Supplementary Fig. 8. Quality control for characterization of CHEK1-CLPB complexes and additional DNA damage control for microscopy.** A, B) Normalized raw intensities and summary of metrics related to the proteins quantified for CHEK1 D130A or L449R AP-MS, respectively. C) Differences in mitochondrial area based on classification as elongated or fragmented. Note that mitochondria were not classified only based on size, but also on shape, branching and other factors. D) Mitochondria of HeLa cells treated with etoposide. The p-value was calculated using a two-tailed t-test. For each condition, 10 images taken across 3 replicates were quantified. All AP-MS experiments were performed in triplicates.

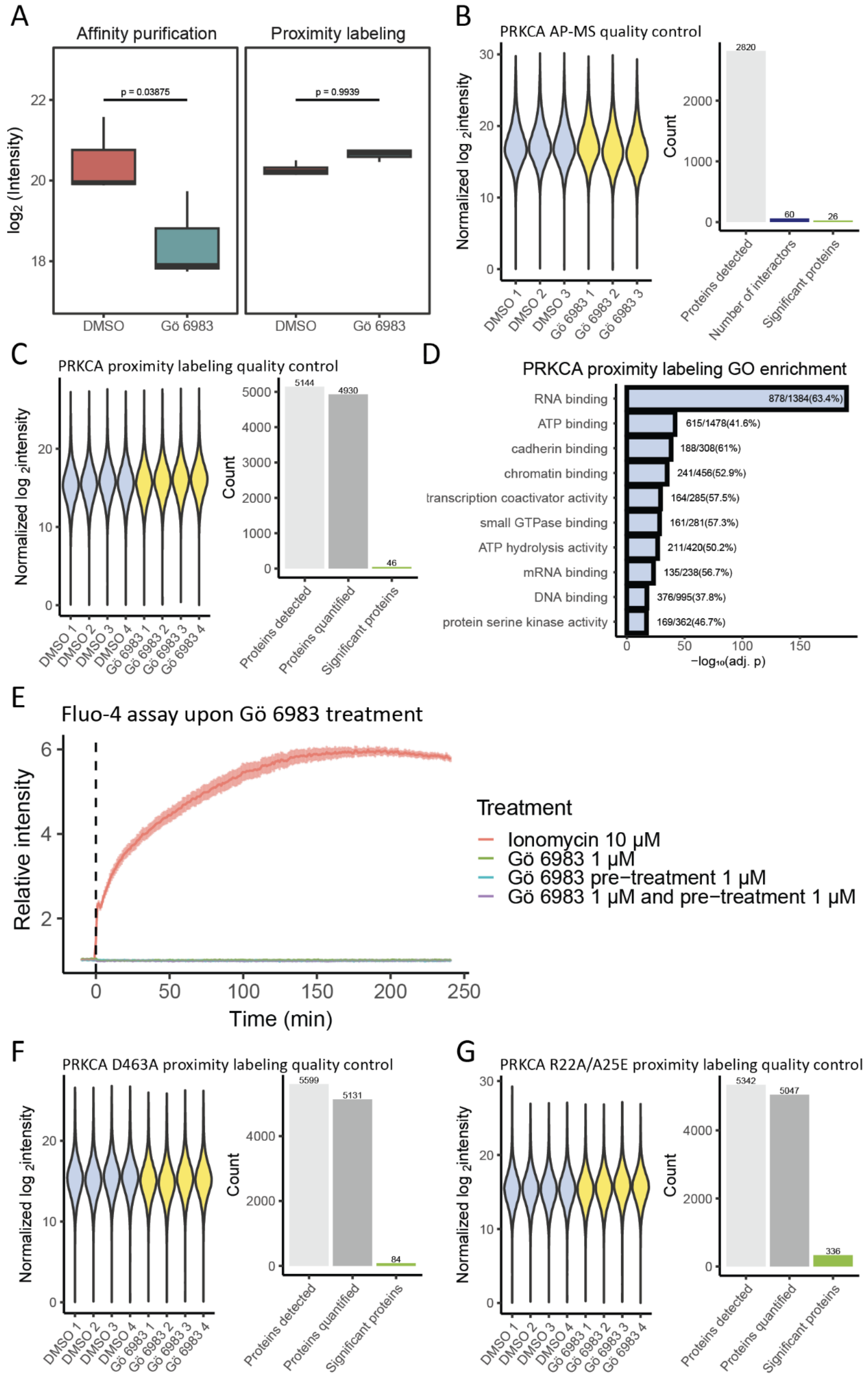


**Supplementary Fig. 9. Quality control of Characterization of the relocalization of PRKCA upon Gö 6983 treatment.** A) Differences in PRKCA abundance upon treatment with Gö 6983 in affinity purification and proximity labeling experiments. The p-value was calculated using a one-tailed t-test. B, C) Normalized raw intensities and summary of metrics related to the proteins quantified for PRKCA AP-MS and *in vivo* proximity labeling, respectively. D) GO enrichment analysis (molecular function) across all proteins quantified in the *in vivo* proximity labeling experiment using PRKCA as bait. Only enriched terms were plotted. P-values were calculated using a Fisher's exact test followed by multiple testing corrections using the procedure of Benjamini-Hochberg. E) We measured changes in intracellular $Ca^{2+}$ concentration in response to Gö 6983 or ionomycin as a positive control using a Fluo-4 calcium imaging assay.

For each time point, the mean and the standard error of the mean are reported. For each condition, six replicates were used. F, G) Normalized raw intensities and summary of metrics related to the proteins quantified for PRKCA D463A and R22A/A25E *in vivo* proximity labeling, respectively. AP-MS experiments were performed in triplicates, and *in vivo* proximity labeling experiments in quadruplicates.

**Supplementary Table 3: List of generated doxycycline-inducible expression vectors and their source.**

| Name | Source |
|---|---|
| N-pTOSH-DCLK1 | Buljan M. *et al.*, *Mol. Cell*, 2020 |
| N-miniTurbo-CAMKK2 | pDONR223-CAMKK2 (addgene 23422) |
| N-pTOSH-CAMKK2 | pDONR223-CAMKK2 (addgene 23422) |
| N-pTOSH-CAMKK2 R311C | Mutated from pDONR223-CAMKK2 |
| N-pTOSH-CAMKK2 D312A | Mutated from pDONR223-CAMKK2 |
| N-pTOSH-CAMKK2 T483D | Mutated from pDONR223-CAMKK2 |
| N-pTOSH-CHEK1 | Buljan M. *et al.*, *Mol. Cell*, 2020 |
| N-pTOSH-CHEK1 D130A | Mutated from N-pTOSH-CHEK1 |
| N-pTOSH-CHEK1 L449R | Mutated from N-pTOSH-CHEK1 |
| N-miniTurbo-CHEK1 | Varjosalo M. *et al.*, *Cell*, 2008 |
| N-pTOSH-PRKCA wt | pDONR-PRKCA (ORFeome 51011@B05) |
| N-miniTurbo-PRKCA wt | pDONR-PRKCA (ORFeome 51011@B05) |
| N-miniTurbo-PRKCA R22A/A25E | Mutated from pDONR-PRKCA |
| N-miniTurbo-PRKCA D463A | Mutated from pDONR-PRKCA |
| N-miniTurbo-PRKAA1 | pDONR223-PRKAA1 (ORFeome 31002@A11) |
| N-miniTurbo-eGFP | pDONR221_EGFP (addgene 25899) |
| N-pTOSH-eGFP | pDONR221_EGFP (addgene 25899) |

**Supplementary Table 4: List of primers used for site directed mutagenesis.**

| Gene | Name | Sequence: (5' to 3') |
|---|---|---|
| CAMKK2 | R311C forward | CTACCAGAAGATCATCCACTGCGACATCAAACCTTCCAACCTCCTGGTCGGAG |
| CAMKK2 | R311C reverse | CTCCGACCAGGAGGTTGGAAGGTTTGATGTCGCAGTGGATGATCTTCTGGTA G |
| CAMKK2 | D312A forward | CCAGAAGATCATCCACCGTGCGATCAAACCTTCCAACCTCCTGGTCGGAGAAG |
| CAMKK2 | D312A reverse | CTTCTCCGACCAGGAGGTTGGAAGGTTTGATCGCACGGTGGATGATCTTCTG G |
| CAMKK2 | T483D forward | CAAACACATTCCCAGCTTGGCAGATGTGATCCTGGTGAAGACC |
| CAMKK2 | T483D reverse | GGTCTTCACCAGGATCACATCTGCCAAGCTGGGAATGTGTTTG |
| CHEK1 | D131A forward | GCATGGTATTGGAATAACTCACAGGGCGATTAAACCAG |
| CHEK1 | D131A reverse | CTGGTTTAATCGCCCTGTGAGTTATTCCAATACCATGC |
| CHEK1 | L449R forward | CATGACGTTATCTAACTTCAGGGCCCTATAAATGATTCCTC |
| CHEK1 | L449R reverse | GGAAGTGTCTCTTGAACTCACGTCCATCACCCTTAGAAAGCCG |
| PRKCA | R22A/A25E forward | CCGCTTCGCCGCCAAAGGGGAGCTGAGGCAG |
| PRKCA | R22A/A25E reverse | CTGCCTCAGCTCCCCTTTGGCGGCGAAGCGG |
| PRKCA | D463A forward | GAGGAATCATTTATAGGGCCCTGAAGTTAGATAACGTCATG |
| PRKCA | D463A reverse | CATGACGTTATCTAACTTCAGGGCCCTATAAATGATTCCTC |

**Supplementary Table 5: Isolation windows used for DIA methods on the Orbitrap Fusion Lumos Tribrid mass spectrometer.**

| Window | Center m/z | z | Isolation Window size (m/z) |
|---|---|---|---|
| 1 | 358 | 2 | 16 |
| 2 | 373 | 2 | 16 |
| 3 | 388 | 2 | 16 |
| 4 | 403 | 2 | 16 |
| 5 | 418 | 2 | 16 |
| 6 | 433 | 2 | 16 |
| 7 | 448 | 2 | 16 |
| 8 | 463 | 2 | 16 |
| 9 | 478 | 2 | 16 |
| 10 | 493 | 2 | 16 |
| 11 | 508 | 2 | 16 |
| 12 | 523 | 2 | 16 |
| 13 | 538 | 2 | 16 |
| 14 | 553 | 2 | 16 |
| 15 | 568 | 2 | 16 |
| 16 | 583 | 2 | 16 |
| 17 | 598 | 2 | 16 |
| 18 | 613 | 2 | 16 |
| 19 | 628 | 2 | 16 |
| 20 | 643 | 2 | 16 |
| 21 | 659 | 2 | 18 |
| 22 | 676 | 2 | 18 |
| 23 | 693 | 2 | 18 |
| 24 | 710 | 2 | 18 |
| 25 | 727 | 2 | 18 |
| 26 | 744 | 2 | 18 |
| 27 | 761 | 2 | 18 |
| 28 | 778 | 2 | 18 |
| 29 | 795 | 2 | 18 |
| 30 | 813 | 2 | 20 |
| 31 | 832 | 2 | 20 |
| 32 | 851 | 2 | 20 |
| 33 | 870 | 2 | 20 |
| 34 | 889 | 2 | 20 |
| 35 | 908 | 2 | 20 |
| 36 | 929.5 | 2 | 25 |
| 37 | 953.5 | 2 | 25 |
| 38 | 977.5 | 2 | 25 |
| 39 | 1006.5 | 2 | 35 |
| 40 | 1048 | 2 | 50 |
| 41 | 1111 | 2 | 78 |

**Supplementary Table 6: Isolation windows used for DIA methods on the Orbitrap Exploris 480 mass spectrometer.**

| Number | Lower bound [m/z] | Upper bound [m/z] | Window size [m/z] |
|---|---|---|---|
| 1 | 350 | 366 | 16 |
| 2 | 365 | 381 | 16 |
| 3 | 380 | 397 | 17 |
| 4 | 396 | 411 | 15 |
| 5 | 410 | 426 | 16 |
| 6 | 425 | 441 | 16 |
| 7 | 440 | 456 | 16 |
| 8 | 455 | 471 | 16 |
| 9 | 470 | 486 | 16 |
| 10 | 485 | 501 | 16 |
| 11 | 500 | 516 | 16 |
| 12 | 515 | 531 | 16 |
| 13 | 530 | 546 | 16 |
| 14 | 545 | 561 | 16 |
| 15 | 560 | 576 | 16 |
| 16 | 575 | 591 | 16 |
| 17 | 590 | 606 | 16 |
| 18 | 605 | 621 | 16 |
| 19 | 620 | 636 | 16 |
| 20 | 635 | 651 | 16 |
| 21 | 650 | 668 | 18 |
| 22 | 667 | 685 | 18 |
| 23 | 684 | 702 | 18 |
| 24 | 701 | 719 | 18 |
| 25 | 718 | 736 | 18 |
| 26 | 735 | 753 | 18 |
| 27 | 752 | 770 | 18 |
| 28 | 769 | 787 | 18 |
| 29 | 786 | 804 | 18 |
| 30 | 803 | 823 | 20 |
| 31 | 822 | 842 | 20 |
| 32 | 841 | 861 | 20 |
| 33 | 860 | 880 | 20 |
| 34 | 879 | 899 | 20 |
| 35 | 898 | 918 | 20 |
| 36 | 917 | 942 | 25 |
| 37 | 941 | 966 | 25 |
| 38 | 965 | 990 | 25 |
| 39 | 989 | 1024 | 35 |
| 40 | 1023 | 1073 | 50 |
| 41 | 1072 | 1150 | 78 |

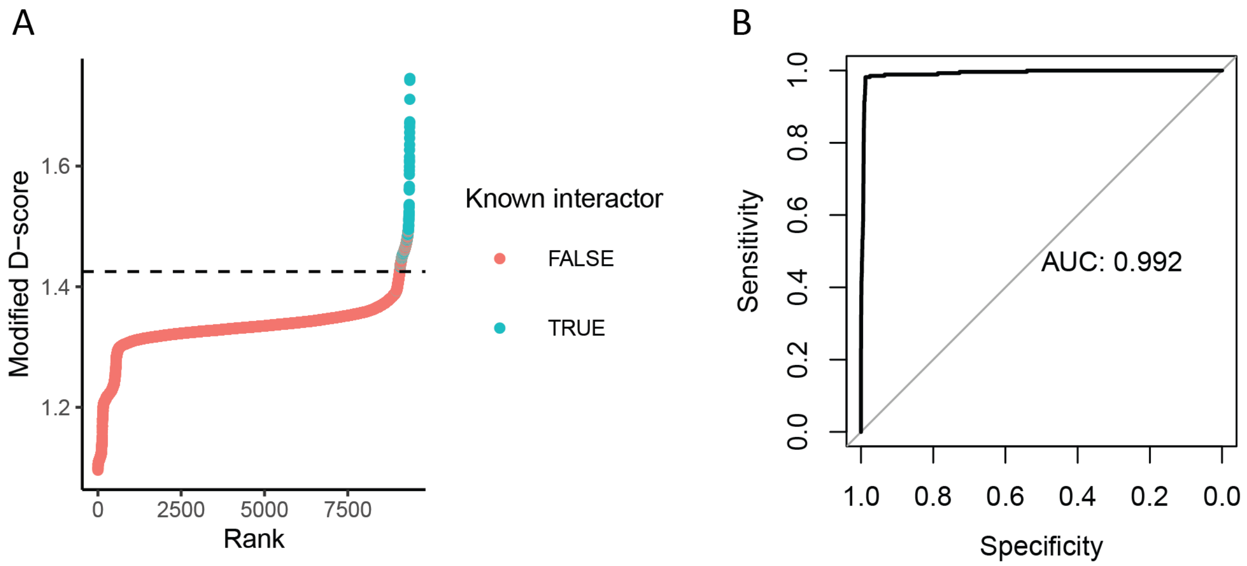


**Supplementary Fig. 10. Determination of interactors using a modified WD score for AP-MS data.** A) Ranking of the modified D-score of known and unknown interactors across all baits. The cutoff was set at 1.425. Any protein above this cutoff in either treatment condition was considered a potential interactor. B) ROC AUC analysis of the modified D-score predicting known interactors across all baits. The AUC is 0.992. The diagonal reference line shows AUC = 0.5.

**Supplementary table 7: Sequences used for the prediction of the CAMKK2-PRKAA1 complex.** Sequences correspond to the kinase domains as annotated on UniProt.

| Gene | Accession | Sequence |
|---|---|---|
| CAMKK2 | Q96RR4 | YTLKDEIGKGSYGVVKLAYNENDNTYYAMKVLSKKKLIRQAGFPRRPPPRGTRPAPGGCIQPRGPIE QVYQEIAILKKLDHPNVVKLVEVLDDPNEDHLYMVFELVNQGPVMEVPTLKPLSEDQARFYFQDLIK GIEYLHYQKIIHRDIKPSNLLVGEDGHIKIADFGVSNEFKGSDALLSNTVGTPAFMAPESLSETRKIFSG KALDVWAMGVTLYCFVFGQCPFMDERIMCLHSKIKSQALEFPDQPDIAEDLKDLITRMLDKNPESRI VVPEIKLHPWV |
| PRKAA1 | Q13131 | YILGDTLGVGTFGKVKVGKHELTGHKVAVKILNRQKIRSLDVVGKIRREIQNLKLFRHPHIIKLYQVIST PSDIFMVMEYVSGGELFDYICKNGRLDEKESRRLFQQILSGVDYCHRHMVVHRDLKPENVLLDAHM NAKIADFGLSNMMSDGEFLRTSCGSPNYAAPEVISGRLYAGPEVDIWSSGVILYALLCGTLPFDDDH VPTLFKKICDGIFYTPQYLNPSVISLLKHMLQVDPMKRATIKDIREHEWF |
